# Impaired hippocampal circuit function underlying memory encoding and consolidation precede robust Aβ deposition in a mouse model of Alzheimer’s disease

**DOI:** 10.1101/2024.05.26.595168

**Authors:** Hanyan Li, Zhuoyang Zhao, Aline Fassini, Han K. Lee, Reese J. Green, Stephen N. Gomperts

**Affiliations:** Department of Neurology, Massachusetts General Hospital, Boston, Massachusetts, USA; Harvard Medical School, Boston, Massachusetts, USA

**Keywords:** Alzheimer’s disease, amyloid-beta, hippocampus, place cell, electrophysiology, sleep

## Abstract

Current therapeutic strategies for Alzheimer’s disease (AD) target amyloid-beta (Aβ) fibrils and high molecular weight protofibrils associated with plaques, but molecular cascades associated with AD may drive neural systems failure before Aβ plaque deposition in AD. Employing hippocampal electrophysiological recordings and dynamic calcium imaging across the sleep-wake cycle in a freely behaving mouse model of AD before Aβ plaques accumulated, we detected marked impairments of hippocampal systems function: In a spatial behavioral task, phase-amplitude coupling (PAC) of the hippocampal theta and gamma oscillations was impaired and place cell calcium fluctuations were hyper-synchronized with the theta oscillation. These changes were not observed in REM sleep. In subsequent slow wave sleep (SWS), place cell reactivation was reduced. These degraded neural functions underlying memory encoding and consolidation support targeting pathological processes of the pre-plaque phase of AD to treat and prevent hippocampal impairments.

## Introduction

In Alzheimer’s disease (AD), deposition of amyloid-beta42 (Aβ) in amyloid plaques and aggregation of intracellular tau in neurofibrillary tangles are associated with neuronal loss and progressive memory impairment on the basis of hippocampal systems failure ^1^. The advent of amyloid-targeted immunotherapies such as Lecanemab (targeting high molecular weight Aβ protofibrils as well as fibrils in plaques) ^2^, Donanemab (targeting the pyroglutamate modification seen in Aβ fibrils in plaques) ^3^, and Aducanumab (targeting Aβ fibrils in plaques) ^4^ for the treatment of AD heralds the beginning of a new era, with evidence to date demonstrating marked reductions of brain amyloid. Even so, these treatments as yet have only modestly improved cognitive outcomes of AD patients ^2,3,5^. Several plausible explanations may explain this discrepancy, including the possibility of problems with the timing or molecular targets of these state-of-the-art treatments against neurodegeneration in AD.

Hippocampal functions are diffusely disrupted in Aβ plaque-laden mice such as the APPswe/PS1dE9 (APP/PS1) transgenic model that harbors genetic mutations that drive production of the Aβ42 peptide and cause autosomal dominant AD ^6,7^. During online memory encoding processes, hippocampal place cells form place fields that are activated sequentially as animals explore an environment in coordination with the hippocampal theta oscillation ^8,9^. In Aβ plaque-laden mice, place representations ^10,11^ and the coupling of the amplitude of gamma oscillations to the phase of theta ^12–19^ are impaired, and neuronal calcium fluctuations hypersynchronize with the theta oscillation ^20^. Aβ plaque-laden mice also show impaired off-line hippocampal memory consolidation processes: In SWS, reduced reactivation of hippocampal place cell pairs ^21^ and reduced coordination of hippocampal sharp wave ripples (SWRs) with cortical spindles and slow oscillations (SO) ^20,22^; in REM sleep, the same oscillatory impairments seen in wakefulness, namely, reduced theta-gamma coupling ^20,23,24^, and hyper-synchronization of neuronal calcium fluctuations with theta ^20^.

Some molecular species arising from aberrant degradation of the amyloid precursor protein (APP) have been found to be cytotoxic ^25^, including soluble Aβ oligomers (AβO). AβO impair synapse function, plasticity, and density ^26–36^ and have been found to drive hippocampal neuronal hyper-excitability in anesthetized AD models ^37–39^. As AβO arise before plaques form, we hypothesized that impairments of hippocampal systems functions underlying memory encoding and consolidation would be evident in advance of plaque deposition. To test this hypothesis, here we acquired hippocampal dynamic calcium imaging and hippocampal and cortical electrophysiological recordings across behavioral states in freely-behaving APP/PS1 transgenic mice in the window before robust plaque deposition ^6,40^. We found marked impairments of hippocampal functions underlying memory encoding and consolidation processes despite minimal Aβ plaque burden.

## Results

To evaluate hippocampal function before plaque deposition, we acquired hippocampal CA1 dynamic calcium imaging with GCaMP6f using the Inscopix miniscope^TM^ together with simultaneous cortical and hippocampal local field potential (LFP) recordings of young APP/PS1 mice (5.2 ± 0.6 months, n=8, male) and their littermate controls (4.7 ± 0.5 months, n=7, male; age, p=0.4, T=0.8, df=13, t-test), as animals explored a linear track (RUN), and subsequently in quiet wakefulness (QW), slow wave sleep (SWS) and REM sleep in their home cage (Fig. 1a). Recording lengths of each state were comparable between genotypes (Fig. S1).

**Figure 1.**
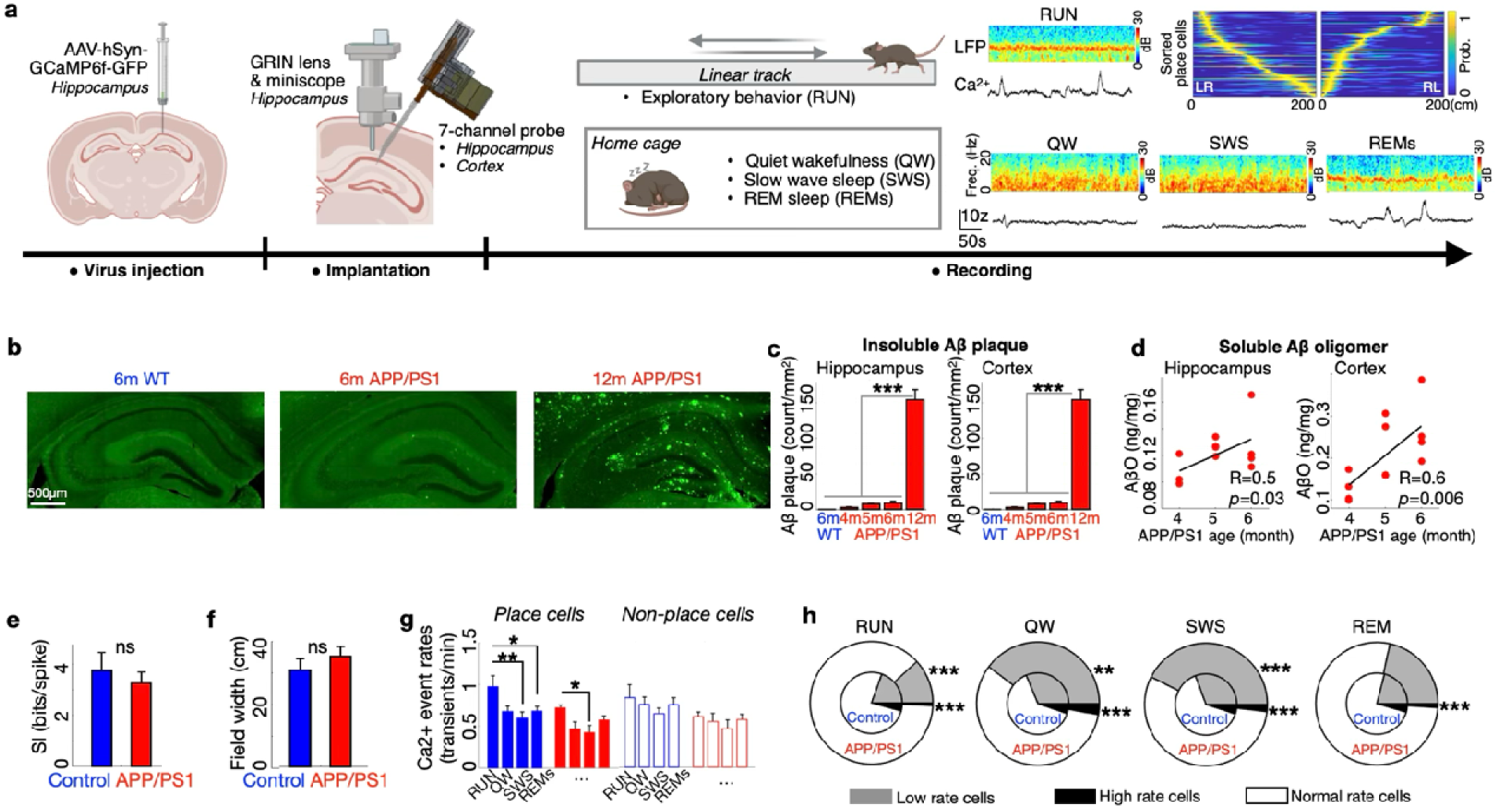
Young APP/PS1 mice expressing minimal Aβ plaque burden and abundant soluble AβO showed preserved place cell function and no calcium hyperactivity across the sleep-wake cycle. a) Schematic of hippocampal calcium imaging and simultaneous ipsilateral LFP recording across the sleep-wake cycle. Insert: examples of LFP spectrogram (color bar indicates power) and one CA1 neuron’s calcium trace across behavioral states; and example dataset of place cells sorted by their place fields (color bar indicates probability) in inbound (LR) and outbound (RL) directions. Yellow shows the place field___of each cell. Exploratory behavior (RUN); quiet wakefulness (QW); slow wave sleep (SWS); rapid eye movement (REM) sleep. **b)** Immunohistochemistry against Aβ in hippocampus of a wild-type control mouse (6 months) and APP/PS1 mice at 6 and 12 months. Green: Aβ plaques (D54D2). **c)** Aβ plaque counts in hippocampus and cortex were low in 4-6 month-old APP/PS1 mice and increased markedly at 12 months. **d)** Levels of soluble AβO in hippocampus and cortex increased with age in APP/PS1 mice. **e)** Spatial information (SI) and **f)** place field width were comparable across genotypes. **g)** Calcium event rates of place cells and non place cells in each behavioral state were similar to those of littermate control mice, and calcium event rates of place cells were higher in RUN than SWS in both genotypes. **h)** In RUN, APP/PS1 had fewer low rate and high rate cells; In QW and SWS, APP/PS1 had more low rate cells and fewer high rate cells. In REM, APP/PS1 had fewer high rate cells. * p<0.05, ** p<0.01, ***p<0.001; c, one-way ANOVA with post-hoc comparisons; d, Pearson correlation; e, f, t-test; g, two-way ANOVA with post-hoc comparisons; h, mixed effects model. b-d: each group n=4 mice; e-h: Control n=5 mice, 395 place cells, 2169 total cells; APP/PS1 n=6 mice, 237 place cells, 2133 total cells. Data are represented as mean ± S.E.M.

### Minimal burden of Aβ plaques and abundant soluble Aβ oligomers in young APP/PS1 mice

In the APP/PS1 model, Aβ plaques have been reported to first deposit around 6 months, with dense accumulation by 12 months ^6,40^. In line with these reports, we observed minimal deposition of Aβ plaques in hippocampus and cortex in young APP/PS1 mice, in contrast to 12-month-old mice (hippocampus: repeated measures ANOVA F(4,15)=42.1, p<0.001, post hoc comparison with positive control p<0.001; cortex: repeated measures ANOVA F(4,15)=105.4, p<0.001, post hoc comparison with positive control p<0.001, Fig. 1b, c). Soluble AβOs were detected in both hippocampus and cortex and increased with age (hippocampus R=0.5, p=0.03; cortex R=0.6, p=0.006, Pearson correlations, Fig. 1d, consistent with prior results ^41,42^).

### Intact place representations but reduced dynamic calcium activity in quiet wakefulness and slow wave sleep before plaque deposition

Behavior was comparable across young APP/PS1 and littermate control groups, with similar run velocity (Control: 4.1 ± 0.7 cm/s; APP/PS1: 3.5 ± 0.6 cm/s, p=0.2, T=1.3, df=13, t-test) and laps on the track (Control: 34.2 ± 3.2 laps; APP/PS1: 39.8 ± 2.5 laps, p=0.2, T=1.3, df=13, t-test), consistent with prior reports ^43^.

To investigate place cell representations across the genotypes, which are often directional ^44^, we considered inbound (LR) and outbound (RL) trajectories separately (Fig. 1a insert). The proportion of place cells was similar in young APP/PS1 and control mice (Control: 16.8% ± 1.9%; APP/PS1: 11.0% ± 2.1%. p=0.07, T=1.9, df=9, t-test). Spatial information (p=0.5, T=0.5, df=9, t-test, Fig. 1e) and place field width (p=0.3, T=0.9, df=9; t-test, Fig. 1f) were also similar across the genotypes. These results are consistent with prior work ^45^.

Previous studies in plaque-laden AD models have detected aberrant neuronal activity ^20,37,38^, and studies in young anesthetized AD mice ^37,39^ have reported hyperactivity of hippocampal neurons before amyloid plaque formation. We therefore extended analyses of neuronal activity of young APP/PS1 mice across the sleep-wake cycle, free from the complex effects of anesthesia ^39^. Compared to littermate control mice, young APP/PS1 mice had similar place cell and nonplace cell mean calcium event rates in each behavioral state (Control place cell vs APP/PS1 place cell: RUN p = 0.07, QW p = 0.1, SWS p = 0.1, REM p = 0.1; Control non place cell vs APP/PS1 non place cell: RUN p = 0.1, QW p = 0.1, SWS p = 0.1, REM p = 0.1; two-way repeated-measures ANOVA with post-hoc comparisons, Fig. 1g, Supplementary Table 1, 2). We did not observe hyperactivity (defined as calcium event rates > 2SD above the control group mean ^20^) in young APP/PS1 mice in any behavioral state (high rate cell proportions, Control vs APP/PS1, RUN: 4.2% ± 0.4% vs 0.7% ± 0.6%, p<0.001; QW: 5.1% ± 0.4% vs 2.8% ± 1.9%, p=0.001; SWS: 4.9% ± 0.3% vs 1.8% ± 1.5%, p<0.001; REM sleep: 4.4% ± 0.5% vs 1.0% ± 0.8%, p<0.001. Mixed effects model, Fig 1h). Instead, we observed an increased proportion of hypoactive cells (< 0.25 spikes/min ^20,37^) in APP/PS1 mice in QW and SWS (low rate cell proportions, Control vs APP/PS1, RUN: 19.5% ± 8.2% vs 11.7% ± 1.5%, p<0.001; QW: 31.2% ± 4.7% vs 39.8% ± 5.8%, p=0.009; SWS: 30.6% ± 3.5% vs 42.5% ± 11.5%, p<0.001; REM sleep: 21.0% ± 4.0% vs 21.5% ± 3.5%, ns. Mixed effects model, Fig 1h), regardless of place cell identity. This finding persisted when high rate cells were defined using a calcium event rate threshold of 3 Z scores: APP/PS1 mice showed no hyperactivity in any behavioral state (RUN p<0.001, QW p=0.001. SWS p<0.001; REM p<0.001, Mixed effects model).

Consistent with prior results in plaque-laden APP/PS1 mice ^20^ (but see ^39^), calcium event rates of place cells were higher in RUN than in SWS in both APP/PS1 and control mice (Control: RUN vs SWS p = 0.009, RUN vs REM p = 0.01; APP/PS1: RUN vs SWS p = 0.02; two-way repeated-measures ANOVA with post-hoc comparisons, Fig. 1g, Supplementary Table 1, 2). In both APP/PS1 and control mice, the calcium event rates of non-place cells were similar across behavioral states.

### Reduced place cell pairwise reactivation despite intact slow wave sleep oscillations before robust plaque deposition

Since place cell properties of young APP/PS1 mice were essentially intact at the single cell level, we next evaluated place cell coordination in offline behavioral states. Place cells with overlapping place fields during spatial behavioral tasks tend to fire together during subsequent rest and sleep in the service of memory consolidation ^46–48^, and prior work has found that reactivation of place cell pairs in SWS is reduced in Aβ plaque-bearing mice ^21^. We therefore set out to evaluate whether place cell reactivation is perturbed prior to robust Aβ plaque deposition. SWR frequency and duration in SWS were comparable in APP/PS1 and control mice (SWR rate: Control 0.061 ± 0.014 event/s, APP/PS1 0.089 ± 0.020 event/s, p=0.2, T=1, df=13; duration: Control 0.024 ± 0.001 s, APP/PS1 0.025 ± 0.001 s, p=0.2, T=1.2, df=13, t-tests). We calculated the spike time cross-correlation of all place cell pairs during exploratory behavior and in subsequent offline states, and identified reactivation when a place cell pair was coactive in both online and offline states ^48^ (Fig. 2a, see Methods and Fig. S2). Pairwise reactivation was detected in each genotype by comparing to the distribution of 500 cross-correlations of shuffled spike times (Supplementary Fig. S3a). The percentage of place cell pairs that reactivated in SWS was significantly reduced in APP/PS1 compared to control mice (p=0.005, two-way repeated-measure ANOVA with post-hoc comparison, Fig. 2a, b, Supplementary Table 3).

**Figure 2.**
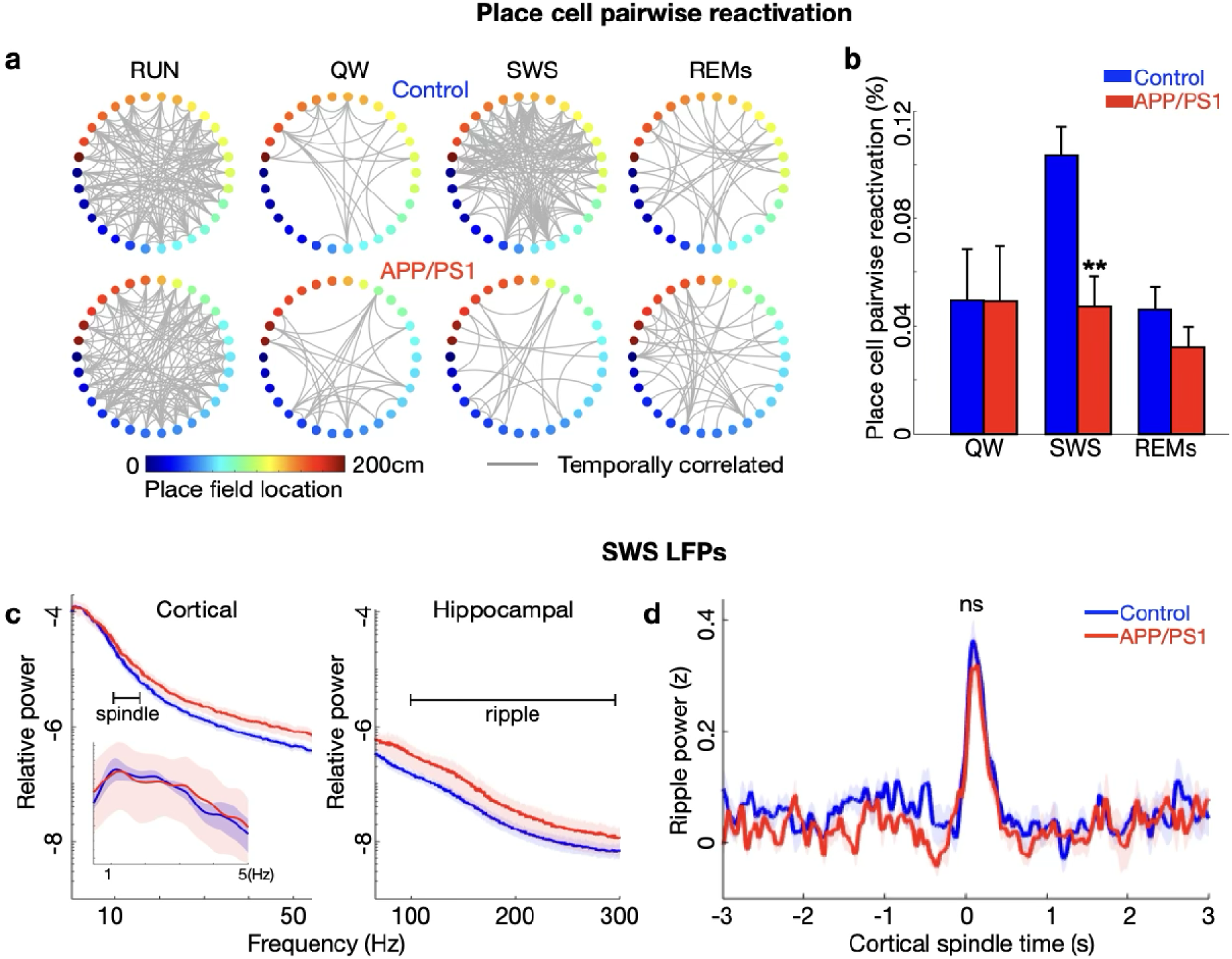
Young APP/PS1 mice showed reduced place cell pairwise reactivation but intact oscillations of slow wave sleep. **a)** Spike time cross-correlations of 30 randomly selected place cells from one control and one APP/PS1 mouse, sorted by place field location on the track (colorbar). Grey line: coactivation. **b)** The percentage of place cell pairs that were temporally reactivated in slow wave sleep (SWS) was reduced in APP/PS1 compared to control mice, but not in quiet wakefulness (QW) or REM sleep (REMs). **c)** In SWS, power in the cortical slow oscillation, delta, spindle (left panel) and hippocampal ripple band (right panel) wa comparable between genotypes. Inset: cortical slow oscillation and delta power. **d)** Cortical spindle-triggered ripple power was comparable between genotypes. **p < 0.01, two-way ANOVA with post-hoc comparison. Control n=5 mice, 395 place cells, 2169 total cells; APP/PS1 n=6 mice, 237 place cells, 2133 total cells. Data are represented as mean ± S.E.M.

Place cell activity and numbers were comparable in APP/PS1 and control mice (see above), and this difference persisted after matching place cell event rate and place cell numbers in control mice and APP/PS1 mice (Supplementary Fig. S3b-d). In contrast to SWS, pairwise reactivation of APP/PS1 and control mice was comparable in QW (as well as in REM sleep) (QW p=0.9, REM p=0.2, two-way repeated-measure ANOVA with post-hoc comparison, Fig 2a, b). Thus, hippocampal reactivation of place cell pairs in SWS was reduced in APP/PS1 mice prior to robust Aβ plaque deposition.

To investigate possible mechanisms underlying reduced APP/PS1 place cell reactivation, we next evaluated the coordination of SWS hippocampal and cortical oscillations. Hippocampal SWRs, cortical spindles, cortical slow oscillations and their nested coordination are fundamental feature of SWS that contribute to memory consolidation ^49^ ^50^. Whereas these SWS oscillations and their coordination are degraded in Aβ plaque-laden AD models ^20,22^, before plaques predominate we found that they remained intact. Specifically, SWR rates and duration (see above), cortical SO power, cortical spindle power, and hippocampal ripple power (SO, p=0.3, T=1, df=13; spindle p=0.1, T=1.4, df=13; ripple: p=0.3, T=1, df=13, t-tests, Fig. 2c) were comparable between young APP/PS1 and control mice, and the coordination of cortical spindles with hippocampal ripples was similar across genotypes (p=0.1, T=1.5, df=13, t-test, Fig. 2d). Thus, in contrast to impaired place cell pairwise reactivation at this early stage of Aβ amyloidosis, the characteristic hippocampal and cortical oscillations of SWS remain intact.

### Impairment of online theta-gamma coupling before plaque deposition

Since the physiological oscillations of SWS were intact in young APP/PS1 mice, we considered the possibility that the reduction in place cell pairwise reactivation could be related to changes in online oscillatory physiology linked to memory encoding. Intra-hippocampal and hippocampal-cortical theta-gamma phase amplitude coupling (PAC) both contribute to memory encoding^13,24^ and degrade in Aβ plaque-bearing AD models ^11,20,51–54^. We found that hippocampal theta and gamma power were comparable across young APP/PS1 and control mice (theta p=0.9, T=0.01, df=13; gamma p=0.4, T=0.8, df=13, t-tests, Fig. 3a), and theta power was not significantly correlated with age in either group (p>0.05, Pearson’s correlation, Fig. 3a). However, intra-hippocampal theta (6-8 Hz) -gamma (40-80 Hz) PAC in APP/PS1 mice was impaired (p=0.03, T=2.3, df=13, t-test, Fig. 3b). Consistent with the possibility of accumulating toxicity, hippocampal theta-gamma PAC in APP/PS1 mice was negatively correlated with age (APP/PS1 R=-0.7, p=0.02; Control p=0.2; Pearson correlations, Fig. 3c). These results were unchanged after controlling for the possible contribution of theta harmonics ^55^ (Supplementary Fig. S4 a-c).

**Figure 3.**
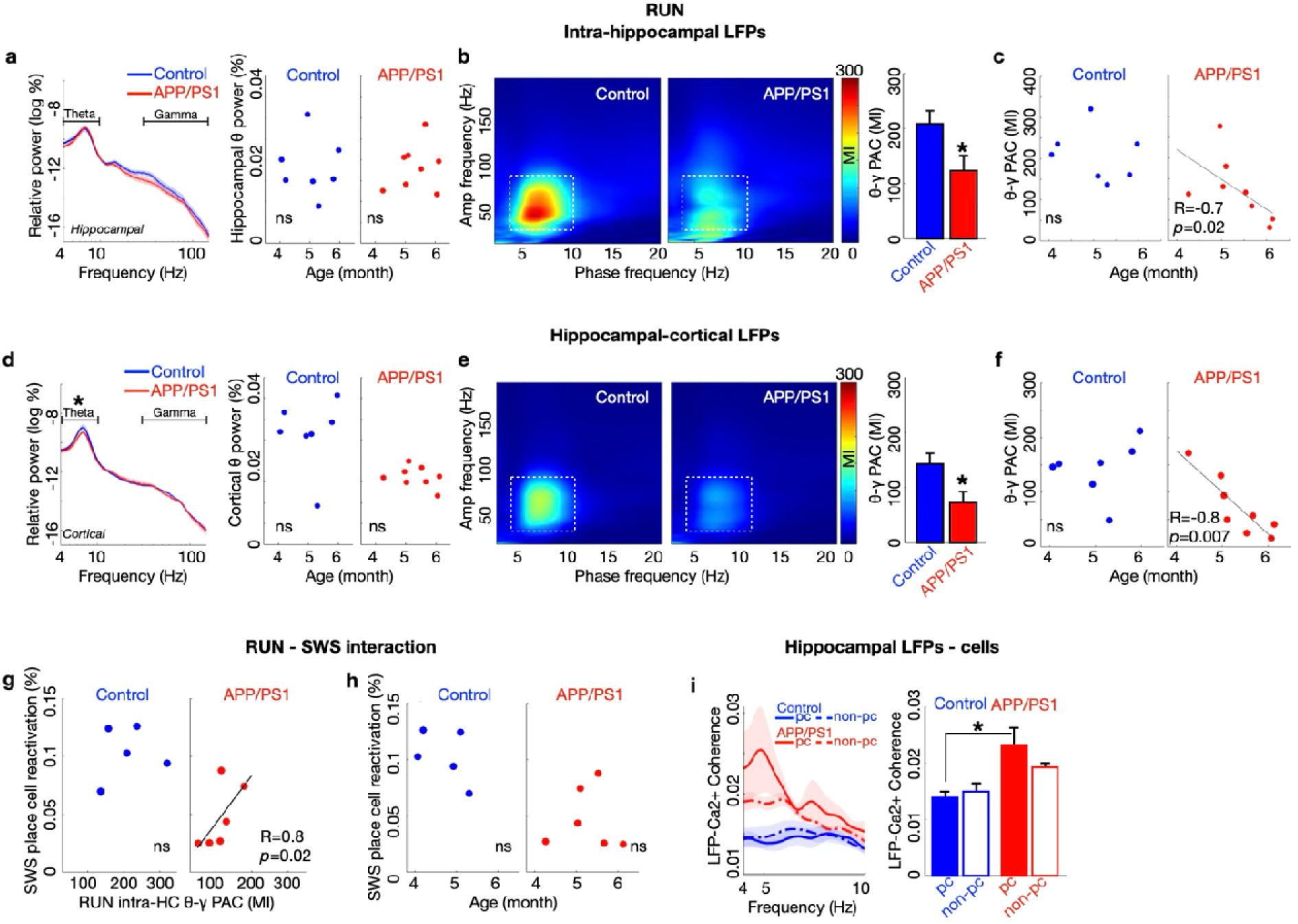
In pre-plaque APP/PS1 mice, theta-gamma phase-amplitude coupling in exploratory behavior was impaired and dynamic calcium fluctuations were excessively synchronized with hippocampal theta. **a)** In RUN, hippocampal theta and gamma power in APP/PS1 and control mice were comparable, and hippocampal theta power was not significantly correlated with age in either group. **b)** Intra-hippocampal theta (6-8Hz) - gamma (40-80Hz) phase - amplitude coupling (PAC) was reduced in APP/PS1 compared to control mice. **c)** Intrahippocampal theta- gamma PAC was negatively correlated with the age of APP/PS1 mice but not control mice. **d)** Cortical theta power was reduced in APP/PS1 mice while cortical gamma power was unchanged. Cortical theta power was not significantly correlated with age in either group. **e)** PAC of hippocampal theta (6-8Hz) - cortical gamma (40-80Hz) was reduced in APP/PS1 compared to control mice. **f)** The reduction of hippocampal theta- cortical gamma PAC was negatively correlated with age in APP/PS1 but not control mice. **g)** In APP/PS1 but not controls, the extent of place cell pairwise reactivation in SWS correlated with the magnitude of intra-hippocampal theta-gamma PAC in RUN. **h)** The proportion of reactivated place cell pairs in SWS was not correlated with age. **i)** Average theta-range LFP-cellular coherence of place cells was higher in APP/PS1 than control mice. *p < 0.05; a, c, d, f, g, h Pearson correlation; a, d, b, e, 2 sided t-test; i, two-way ANOVA with post-hoc comparisons. a – f: Control n=7 mice, APP/PS1 n=8 mice; g – i: Control n=5 mice, 395 place cells, 1774 non-place cells, 2169 total cells; APP/PS1 n=6 mice, 237 place cells, 1896 non-place cells, 2133 total cells. Data are represented as mean ± S.E.M. pc, place cells. Non-pc, nonplace cells.

Cortical gamma power was also comparable between genotypes (p=0.6, T=0.4, df=13, t-test, Fig. 3d), while theta sampled in cortex was reduced in APP/PS1 mice (p=0.005, T=3.3, df=13, t-test; (Fig. 3d). Cortical theta power was not significantly correlated with age in APP/PS1 or control mice (Fig. 3d). In this context, hippocampal theta (6-8 Hz) - cortical gamma (40-80 Hz) PAC was also impaired during exploratory behavior in young APP/PS1 mice (p=0.02, T=2.5, df=13, t-test, Fig. 3e). As with intrahippocampal theta-gamma PAC, in APP/PS1 but not control mice, hippocampal-cortical PAC was negatively correlated with age (APP/PS1 R=-0.8, p=0.007; Control p=0.9; Pearson correlation, Fig. 3f). This result persisted after controlling for the possible contribution of theta harmonics ^12^ (Supplementary Fig. S4 d-f). Together, these results suggest that accumulating APP degradation products impair theta-gamma PAC well before plaques robustly deposit.

Interestingly, the magnitude of hippocampal theta-gamma PAC in RUN was strongly correlated with the extent of place cell pairwise reactivation in subsequent SWS in APP/PS1 mice (APP/PS1 R=0.8, p=0.02; Control p=0.3; Pearson correlations, Fig. 3g). In contrast, pairwise reactivation was not correlated with age (APP/PS1 p=0.2; Control p=0.2; Pearson correlations, Fig. 3h). These observations raise the possibility that the reduction of theta-gamma PAC during online behavior may contribute to the impairment of place cell pairwise reactivation that arises in subsequent SWS.

### Aberrant synchrony of somatic calcium fluctuations with hippocampal theta before plaque deposition

As oscillations both coordinate and reflect the coordination of neuronal activity, and as calcium signaling links synaptic activity to synaptic plasticity and gene expression, we considered the possibility that breakdown in the coordination of neuronal activity reflected in impaired theta-gamma PAC might manifest at the cellular level in altered coordination of somatic calcium fluctuations with the LFP. In exploratory behavior, neuronal calcium levels can synchronize with theta oscillations, and this synchronization is aberrantly increased in Aβ plaque-laden mice ^20^. We therefore measured the coherence of hippocampal neuronal calcium traces with the LFP in APP/PS1 mice before significant plaque accumulation. Compared to control mice, APP/PS1 mice showed hyper-synchronization of place cell calcium fluctuations with theta (p=0.01, two-way repeated-measured ANOVA post-hoc comparisons, Fig. 3i, Supplementary Table 4). Thus, at the pre-plaque stage preceding robust Aβ plaque deposition, dysfunction of memory encoding processes manifest at both the oscillatory and the cellular level.

### REM sleep theta-gamma coupling before plaque deposition

As intra-hippocampal and hippocampal-cortical theta-gamma PAC in REM sleep contribute to memory consolidation ^56,57^ and are diminished in Aβ plaque-bearing AD mice ^20,53^, we evaluated REM sleep theta-gamma PAC prior to robust Aβ plaque deposition. Hippocampal REM sleep theta and gamma power (theta p=0.7, T=0.3, df=13; gamma p=0.5, T=0.6, df=13 t-tests, Fig. 4a), and intra-hippocampal theta (6-8 Hz) -gamma (40-80 Hz) PAC were comparable across genotypes (p=0.9, T=0.01, df=13, t-test, Fig. 4b), consistent with prior results ^53^. In contrast to RUN, hippocampal theta power in REM sleep of APP/PS1 mice was negatively correlated with age (R=-0.8, p=0.008, Pearson correlation, Fig. 4a). This correlation was not observed in control mice. Only a trend level correlation was observed between intra-hippocampal PAC magnitude in REM sleep of APP/PS1 mice and age (R=-0.6, p=0.05, Pearson correlation, Fig. 4c), and this result did not survive after controlling for the possible contribution of theta harmonics (Supplementary Fig. S5 a-c).

**Figure 4.**
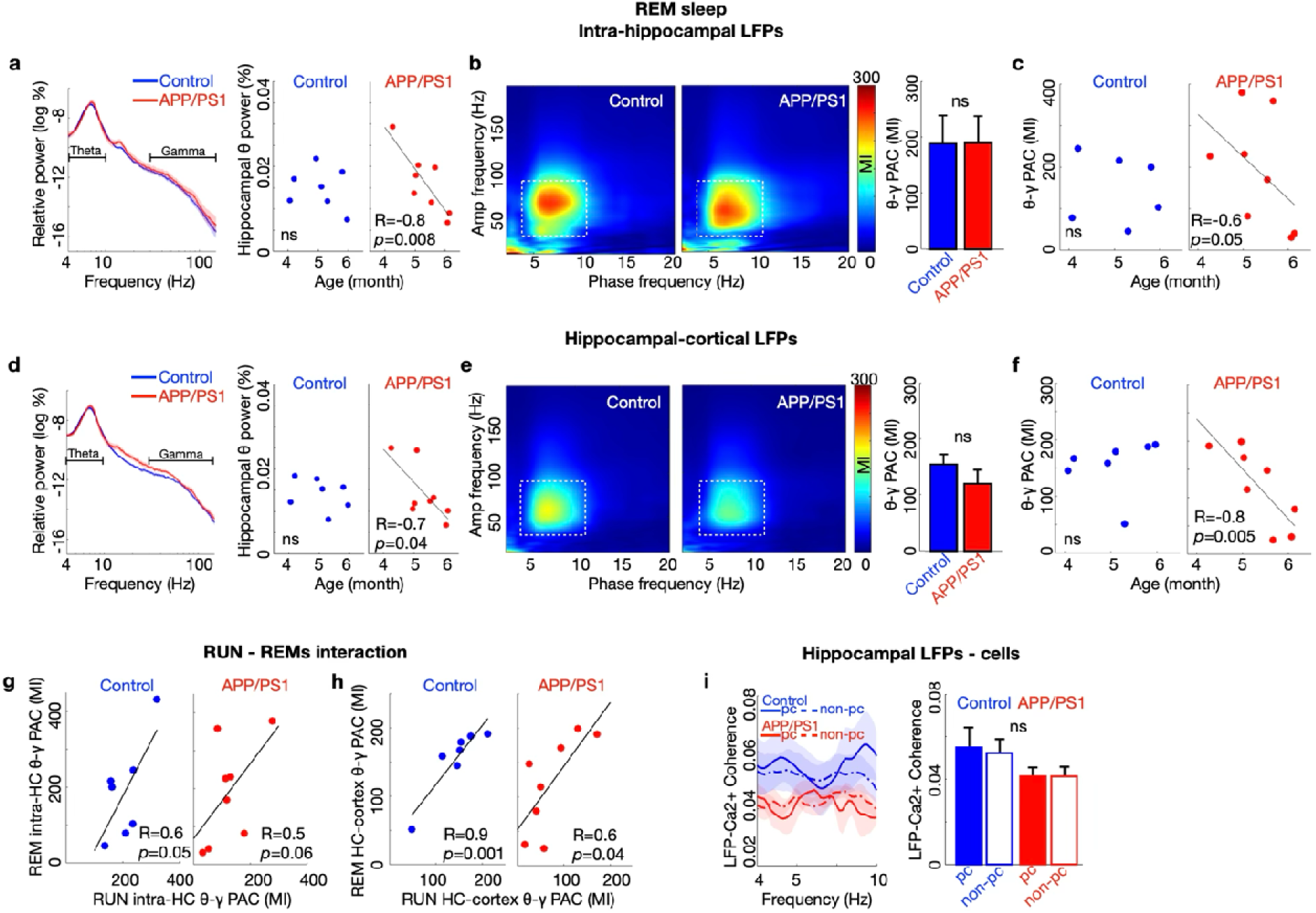
In young APP/PS1 mice, theta-gamma phase amplitude coupling in REM sleep wa intact and correlated with theta-gamma phase-amplitude coupling during exploratory behavior. **a**) In REM sleep, hippocampal theta and gamma power were comparable between groups. Hippocampal theta power of APP/PS1 but not control mice was negatively correlated with age. **b)** Intra-hippocampal theta (6-8Hz) - gamma (40-80Hz) phase-amplitude coupling (PAC) was comparable in APP/PS1 and control mice. **c)** Intrahippocampal theta-gamma PAC showed only a trending negative correlation with the age of APP/PS1 mice but not control mice. **d)** Cortical theta and gamma power were comparable between groups. Cortical theta power of APP/PS1 mice but not control mice was negatively correlated with age. **e)** PAC of hippocampal theta (6-8Hz) - cortical gamma (40-80Hz) was comparable across genotypes. **f)** Hippocampal theta- cortical gamma PAC was negatively correlated with age in APP/PS1 mice but not control mice. **g)** Intra-hippocampal theta-gamma PAC between RUN and REM sleep showed a trending correlation in both genotypes. **h)** Hippocampal theta - cortical gamma PAC in RUN and in REM sleep were significantly correlated in both AP/PS1 and control mice. **i)** Average theta-range LFP-cellular coherence in REM sleep was comparable across APP/PS1 and control mice. a, c, d, f, g, h, Pearson correlation; a, d, b, e, 2 sided t-test; i, two-way ANOVA with post-hoc comparisons. a – g: Control n=7 mice, APP/PS1 n=8 mice; i: Control n=5 mice, 395 place cells, 2169 total cells; APP/PS1 n=6 mice, 237 place cells, 2133 total cells. Data are represented as mean ± S.E.M. pc, place cells. Non-pc, nonplace cells.

REM sleep cortical gamma power (p=0.1, T=1.5, df=13, t-test, Fig. 4d), and hippocampal theta (6-8 Hz) - cortical gamma (40-80 Hz) PAC were also comparable across genotypes (p=0.2, T=1, df=13, t-test, Fig. 4e). Even so, in young APP/PS1 but not control mice, cortical theta power (R=-0.7, p=0.04, Pearson correlation, Fig. 4d) and hippocampal-cortical PAC in REM sleep (R=- 0.8, p=0.005, Pearson correlation, Fig. 4f) were significantly and negatively correlated with age, and this result persisted after controlling for the possible contribution of theta harmonics (Supplementary Fig. S5 d-f). These results suggest that as in exploratory behavior, theta-gamma PAC during REM sleep also falls with accumulation of toxic species before robust plaque deposition.

Because theta-gamma PAC in APP/PS1 mice was differentially affected in RUN and REM sleep but negatively correlated with age in both behavioral states, we compared theta-gamma PAC in RUN versus REM sleep. In both APP/PS1 and control mice, the magnitude of hippocampal theta - cortical gamma PAC in RUN and REM sleep were strongly correlated (Control R=0.9, p=0.001; APP/PS1 R=0.6, p=0.04; Pearson correlations, Fig 4h), with only a trend level correlation between intra-hippocampal theta-gamma PAC in RUN and REM sleep (Control R=0.6, p=0.05; APP/PS1 R=0.5, p=0.06; Pearson correlations, Fig. 4g).

We next sought to determine whether the hyper-synchronization of somatic calcium levels with the theta oscillation that we observed in exploratory behavior (Fig. 3i) extended to REM sleep. In contrast to findings in exploratory behavior, neuronal calcium synchronization with theta in REM sleep was similar across APP/PS1 and control mice (p=0.2, two-way repeated measures ANOVA post-hoc comparison, Fig. 4i, Supplementary Table 5). These results suggest that hippocampal function in REM sleep may be more resilient than in other brain states to the accumulation of toxic APP cleavage products.

## Discussion

In the context of well-described impairments of hippocampal neurophysiology in Aβ plaque-laden mice yet limited therapeutic benefits in manifest AD from targeting Aβ plaques, we investigated hippocampal functions that contribute to memory encoding and consolidation in young APP/PS1 mice harboring AD-associated gene mutations but lacking significant Aβ plaque burden. We found reduced theta-gamma PAC and excessive theta synchronization of hippocampal place cells with their surrounding LFP during exploratory behavior, and reduced place cell pairwise reactivation in subsequent SWS. These diverse impairments of neural functions underlying memory encoding and consolidation manifest before significant plaque burden and implicate other toxic processes in AD.

Hippocampal place cells encode the environment during online behavior, and their network reactivation during offline behaviors, including hippocampal replay during sharp wave ripples ^46,58^, contributes to sleep-dependent learning ^46,59^ and memory consolidation ^47,60,61^. Prior work has reported reduced place cell pairwise reactivation in Aβ plaque bearing mice ^21^, and the present results show that this result arises prior to significant Aβ plaque burden. The impairment of pairwise reactivation did not result from differences in behavior, LFP properties such as ripple rate and duration, or cellular properties such as calcium event rate between young APP/PS1 and control mice. Likewise, although the coordination of cortical slow oscillations, corticothalamic spindles, and hippocampal sharp wave ripples is a key feature of memory consolidation in SWS ^49,50^, and this coordination degrades in Aβ plaque-bearing animals ^20,22^, these oscillations and their coordination were preserved in young APP/PS1 mice lacking robust plaque deposits. As such, the reduced place cell pairwise reactivation observed in these mice stands in contrast with intact SWS oscillatory coordination.

Brain oscillations and their coupling play a crucial role in hippocampal-dependent memory functions. Online intra-hippocampal and hippocampal-cortical theta-gamma PACs have been linked to memory encoding ^12–19^, and, like the coordinated oscillations of SWS, are also impaired in multiple AD models harboring Aβ plaques ^11,20,51–54^. In contrast to the preserved oscillations of SWS, online hippocampal and hippocampal-cortical theta-gamma PACs were reduced in young APP/PS1 mice. As the extent of these reductions correlated with age, online theta-gamma PAC appears to degrade as APP degradation products such as soluble AβO accumulate.

The intra-hippocampal and hippocampal-cortical theta-gamma PACs of REM sleep have also been linked to memory consolidation ^12,13^ . While these deteriorate in plaque-laden AD model mice ^20^, the present findings show that before significant plaque deposition, theta-gamma PACs of REM sleep remain grossly preserved, consistent with prior results ^22^. Even so, a subtle vulnerability was evident, in that hippocampal-cortical theta-gamma PAC declined with increasing age.

In plaque-laden APP/PS1 mice, CA1 neuronal calcium fluctuations are excessively synchronized with hippocampal theta ^20^. Consistent with both this result and the greater vulnerability of online theta-gamma PAC compared to REM sleep in young APP/PS1 mice, hyper-synchronization of neuronal calcium oscillations with the hippocampal LFP was detected in young APP/PS1 mice in exploratory behavior alone. Although the mechanisms underlying impaired theta-gamma PAC and aberrant synchronization of neuronal calcium levels with the theta oscillation remain to be determined, brain oscillations and their coupling are known to reflect the coordinated activity of large populations of excitatory and inhibitory neurons, and their imbalance or dyscoordination may contribute. In this context, it is possible that Aβ-mediated dysfunction of inhibitory neurons and their synapses may contribute to these findings ^62^.

Interestingly, despite the relatively small dataset, the degree of impairment of hippocampal theta-gamma PAC in exploratory behavior was correlated with the reduction of place cell pairwise reactivation in subsequent SWS. This observation raises the possibility of a primary deleterious effect of AβO or other APP degradation product on memory encoding online that then impairs memory consolidation in SWS. Linked impairments across behavioral states mediated by the effects of impaired theta-gamma PAC and hyper-synchronized neuronal calcium fluctuations on theta sequences that link the activity of adjacent place cells into trajectories within each theta cycle is one potential mechanism for this observation ^63–65^. This possibility will be worth assessing in future studies.

Although we observed marked abnormalities of hippocampal function across the sleep-wake cycle of young APP/PS1 mice, we did not detect aberrant neuronal hyperactivity, as has been reported in studies of anesthetized APP/PS1 mice ^37–39^. Instead, we observed a tendency towards hypoactivity in non-theta offline states. Reports of changes in neuronal activity in APP/PS1 mice have been heterogeneous ^39^, and prior work has shown that hyperactivity in young APP/PS1 mice is brought out by and depends on anesthesia and recorded cell types (based on the use of the hsyn vs CAMKII promoter) ^39^. In this context, the present results suggest that hippocampal dynamic calcium hyperactivity in the behaving animal does not appear to be a prominent and requisite finding of early Aβ pathology.

Interestingly, the hippocampal systems level impairments observed here in young APP/PS1 mice precede many of the behavioral impairments that arise in this AD model on standard behavioral assays, although deficits of contextual memory have been reported to start around 6 months ^66,67^. Thus, sampling hippocampal systems level function appears to provide a sensitive readout of brain dysfunction in AD models.

While tau fibrils and neuronal loss are strong predictors of cognitive impairment in AD ^68,69^, the current findings support the early contribution of additional AD-associated processes linked to APP processing. Prior to plaque deposition, APP cleavage products including toxic soluble species of Aβ accumulate. Among these, AβO are well-known to be cytotoxic and have been shown to induce cognitive impairment ^31,36,70^. Among soluble Aβ species, small oligomers are particularly toxic ^36^, and levels of small brain AβO but not monomer have been shown to correlate with AD progression in APP/PS1 mice in the age range studied here ^42^.

Given the cellular and synaptic toxicity of AβO and their relevance to AD progression, as well as the minimal levels of Aβ protofibrils detectable in young APP/PS1 mice ^42^, the accumulation of AβO is well-positioned to contribute to the marked systems-level hippocampal dysfunctions observed here. Even so, other non-Aβ mechanisms resulting from APP overexpression and the PS1 mutation ^25,71^ may also contribute. Among these, non-Aβ APP cleavage products such as sAPP11, APP intracellular domain (AICD) and P3 fragment are biologically active ^72,73^, and soluble APP products have been associated with cognitive impairment, neuronal loss, electrophysiological abnormalities and GABA reduction ^74^. In addition, PS1 mutations through their impact on γ-secretase function affect digestion of γ-secretase substrates including but not limited to APP ^75–77^. Like the PS1dE9 mutation present in the APP/PS1 mouse ^78^, PS1 mutations typically reduce γ-secretase activity, leading to complex downstream effects including altered calcium homeostasis ^79^, increased inflammatory responses in the brain ^76^, and contributing to progressive brain atrophy, memory deficits, and synaptic impairments ^76^. Future experiments to dissect the contributions of each potential mechanism to hippocampal systems failure will be of value.

Strengths and innovations of this study include the use of a well-characterized, freely behaving AD model to evaluate hippocampal functions underlying memory encoding and consolidation before Aβ plaques predominate, free from the complexities associated with anesthesia. The acquisitions of simultaneous hippocampal dynamic calcium imaging and ipsilaterally placed local electrophysiological recordings is a further strength, enabling us to relate brain oscillations critical for normal hippocampal mnemonic functions to neuronal calcium fluctuations, place cell activity, and their reactivation in SWS. Limitations of this study include the use of a single model of amyloidosis, and additional work in other models will be informative. A further limitation is the correlative nature of the work. In this regard, future, causal experiments will prove useful to identify the factors underlying the hippocampal physiological impairments observed here, which could derive from elevated levels of soluble Aβ42 oligomers or other fragments, and/or from presenilin 1 dysfunction.

In summary, the results of this study in AD model mice prior to robust Aβ plaque deposition reveal impairments in hippocampal cellular and oscillatory functions that contribute to both memory encoding and consolidation. Targeting APP related cascades rather than Aβ plaques, per se, may improve upon the significant but, to date, only modest effectiveness of anti-amyloid antibody therapies for AD ^1–4^.

## Materials and Methods

All procedures were approved by the Institutional Animal Care and Use Committees of the Massachusetts General Hospital and the study was conducted in accordance with the ethical guidelines of the US National Institutes of Health. This study is reported in accordance with ARRIVE guidelines.

### Surgical preparation

All procedures were similar to our previous studies ^20,80^. The mouse strain used for this research project, B6C3-Tg(APPswe,PSEN1dE9)85Dbo/Mmjax, RRID:MMRRC_034829-JAX, was obtained from the Mutant Mouse Resource and Research Center (MMRRC) at The Jackson Laboratory, an NIH-funded strain repository, and was donated to the MMRRC by David Borchelt, Ph.D., McKnight Brain Institute, University of Florida ^7^. Briefly, APP/PS1 mice and their age/gender matched non-transgenic control litters were first injected with 1 µL AAV-hSyn-GCaMP6f-GFP into the hippocampal CA1 region (AP −2.1 mm, ML −1.65 mm, DV −1.4 mm) under anesthesia (isoflurane 1.5-2%). Two weeks later, a grin lens (1 mm diameter, AP −2.1 mm, ML −1.65 mm, DV −1.4 mm) and a custom drivable probe holding 7 stereotrodes (AP −2.8 mm, ML 2.6 mm, −36 ° relative to the AP axis, −39 ° relative to the inter-aural axis; DV −1.2 mm) were implanted ipsilaterally, to sample the LFP in the vicinity of the calcium imaging site, with cerebellar ground. The microscope baseplate was attached three weeks later. Carprofen (10 mg/kg, s.c) was used for postoperative analgesia. Animals were housed in individual cages with a 12 hr light/12hr dark-standard light cycle. All mice were male to avoid sex differences in Aβ pathogenesis in APP/PS1 mice ^81^.

### Calcium imaging and electrophysiology recordings

The Inscopix^TM^ nVista3 mini-microscope and acquisition system were used to sample calcium imaging at 20 Hz. Local field potential (LFP) (Neuralynx^TM^) was sampled at 2 kHz followed by 0.5-900 Hz filtering. Animal position was recorded with overhead camera tracking of two LED diodes mounted on the headstage. Electrophysiology and calcium imaging were acquired simultaneously: Electrophysiology was recorded continuously, while calcium imaging was recorded in 10 min acquisition blocks, with 4-5 minute intervals off-camera to avoid photobleaching. These datasets were synchronized and aligned by TTL pulses offline.

### Behavior and sleep stages

Data were acquired as animals first traversed a linear track and were then moved to their home cage within the same recording room for immediate acquisition of subsequent Quiet wakefulness (QW), slow-wave sleep (SWS), and REM sleep. Running behavior on the track was identified when run speed exceeded 1 cm/sec; QW was defined as wakeful immobility (< 0.5 cm/sec, eyes open) in the home cage; SWS was identified when SWRs were present and delta/theta LFP power ratio exceeded 2, while the animal maintained a sleep posture; REM sleep was identified when the theta/delta LFP power ratio exceeded 2 in sleep.

### Data analysis

All analyses were performed using MATLAB (2019b) and were similar to our previous studies ^20,80^.

#### Calcium imaging

Inscopix Data Processing Software (IDPS, Inscopix, Palo Alto, California) was used for movie preprocessing, motion correction, normalization ΔF/F and constrained non-negative matrix factorization (CNMFE) cell identification^82^. The shape and signal-noise-ratio (SNR) of each cell soma was reviewed manually for quality control. Calcium events were then detected at 2 Z scores above the mean, with onset at 30% of peak amplitude. Low rate cells were defined as < 0.25 transients/min ^20,83^; high rate cells were defined using a calcium event rate threshold of 2 Z scores above the mean of the control group for each behavioral state ^20^.

#### Place cell analyses

Spatial tuning curves (3 cm bins) were constructed for each cell for each running direction using all detected calcium events acquired during running. Spatial information was computed and compared to 1000 shuffled spatial information distributions to test significance (Monte Carlo p-value < 0.05) ^84^. Place field width was defined at 50% of peak firing distribution over track location ^85^. Spike time cross-correlations of all place cell pairs in both online (exploratory/RUN) and subsequent offline (QW, SWS) states were calculated. Reactivation was defined when the same pair was coactivated during both online and specific offline states ^48^. Coactivation was defined based on the distinguished clusters (cluster 1 not coactivated: <0.001, and cluster 2 coactivated: >0.02) in the distribution of correlations of all place cell pairs in each state (Fig. S2). Results persisted across a broad range of thresholds (between >0.001 and <0.02), here we used >0.002 per previous work ^48^. Proportions of reactivated pairs among all pairs of each animal were used to compare between genotypes. Recordings with at least 10 place cells (5 out of 7 control mice and 6 out of 8 APP/PS1 mice) were included in the pairwise reactivation analysis.

#### LFP analyses

LFPs were filtered to obtain slow oscillations (0.5-1 Hz), delta (1-4 Hz), theta (4-10 Hz), gamma (30-150 Hz), and ripples (100-300 Hz). The spectrograms of each channel were calculated using the Chronux toolbox ^86^ with a moving window of size 2 s and step size 1 s. A stereotrode in s. radiatum showing high amplitude theta was used for assessments of RUN and REM sleep; a stereotrode in s. pyramidale showing high amplitude SWRs was used for assessments of QW and SWS; a stereotrode within cortex showing slow oscillations and sleep spindles was also used. Power spectra were computed from 0 to 200 Hz using the Chronux toolbox ^86^, and the sum of relative power of each frequency band was used for quantification. Phase-amplitude-coupling (PAC) modulation index (MI) was calculated with 1-25 Hz for phase frequency and 1-300 Hz for amplitude frequency using the MATLAB toolbox https://data.mrc.ox.ac.uk/data-set/matlab-toolbox-estimating-phase-amplitude-coupling ^87,88^, and the mean value of the high coupling region was used for quantification ^22^. Spindle events were detected at 1.25 Z score with minimum length of 200 ms; SWR events were detected at 3 Z score with minimum length of 20 ms ^49^. Ripple power triggered on spindle onsets was averaged ± 0.5 s around lag = 0 s and compared between groups. LFPs were downsampled to the same sampling rate as calcium imaging for the Chronux coherence analysis ^86^ for oscillatory-cellular synchronization ^20^.

### Biochemistry

#### Euthanasia

Mice were euthanized by ketamine-xylazine overdose (240-300 mg/kg ketamine + 15-30 mg/kg xylazine IP).

#### Brain protein extraction

Mice were perfused with cold PBS containing protease inhibitor (cat # A32957), and the brain was rapidly excised. One hemisphere was separated for immunohistochemistry analyses. Cortex and hippocampus were dissected from the remaining hemisphere and immediately frozen in liquid nitrogen, then stored at −80°C before use. Protein extraction was performed as indicated by the ELISA kit. In summary, frozen brain samples were homogenized in 10 vol (w/v) of cold TBS containing protease inhibitor using a Teflon-glass and centrifuged at 16,000 x g for 30 min at 4°C. The pellets were re-suspended in TBS plus 1% Triton X-100 containing protease inhibitor (TBS-T) and centrifuged at 16,000 x g for another 30 min at 4°C. The TBS-T fraction was used to measure oligomeric β-amyloid. The total protein concentration in TBS-T fractions was measured using the BCA protein assay kit (cat# 23225).

#### ELISA

ELISA was used to quantify oligomeric β-amyloid in cortical and hippocampal samples (Biosensis cat# BEK-2215-1P/2P). Oligomeric β-amyloid concentration was determined using a standard curve, and values are reported as ng/mg of protein. Analyses were blinded to animal age.

#### Immunohistochemistry

Hemibrains were drop-fixed in 4% paraformaldehyde for two days and sectioned at 40 μm on a vibratome (Leica, Buffalo Grove, IL). Sections were collected in a 1 in 4 series and stored in PBS at 4°C. Sections were mounted, and antigen retrieval was performed with citrate buffer at 95°C for 30 min. Next, sections were blocked and permeabilized in TBS-T with 5% bovine serum albumin (BSA) and 5% normal goat serum (NGS) for 1h. Sections were incubated overnight in Alexa Fluor 488 Conjugate primary β-Amyloid antibody (D54D2, cat# 51374S) at 4C. Following several washes in PBS, slides were cover slipped with anti-fade mounting media, and images were acquired using an Olympus VS120 Slide Scanner.

### Statistical analysis

All group values were represented as mean ± s.e.m. For analyses of dynamic calcium imaging hypo- and hyperactivity, mixed effects models were used. For the remaining analyses, two-sided *t*-test and two-way ANOVA with repeated measures followed by *post hoc* Tukey-Kramer tests were used for statistical analyses according to the experimental design (SPSS). The significance threshold was set at *p* < 0.05.

## Acknowledgements

We thank Joseph J. Locascio for statistical consultation.

## Author contributions

Conceptualization, H.L., Z.Z. and S.N.G.; Methodology, H.L., Z.Z., A.F. and S.N.G.; Software, H.L. and S.N.G.; Validation, H.L., Z.Z., A.F., H.K.L. and R.J.G.; Formal Analysis, H.L.; Investigation, Z.Z., A.F., H.K.L. and R.J.G.; Resources, H.L., Z.Z., A.F., H.K.L. and R.J.G.; Data Curation, H.L. and R.J.G.; Writing – Original Draft, H.L. and S.N.G.; Writing – Review & Editing, H.L., Z.Z., A.F., H.K.L., R.J.G. and S.N.G.; Visualization, H.L.; Supervision, S.N.G.; Project Administration, S.N.G.; Funding Acquisition, S.N.G.

## Data availability statement

The datasets generated and analyzed in the current study are available in the Dryad repository, https://datadryad.org/stash/share/gN7A2L9rPMErDH74jxAWQUdjKm7A89v4ERaftNsJOiA

## References

1. Bloom, G. S. Amyloid-β and Tau: The Trigger and Bullet in Alzheimer Disease Pathogenesis. JAMA Neurology 71, 505–508 (2014).

2. van Dyck, C. H. et al. Lecanemab in Early Alzheimer’s Disease. New England Journal of Medicine 388, 9–21 (2023).

3. Mintun, M. A. et al. Donanemab in Early Alzheimer’s Disease. N Engl J Med 384, 1691– 1704 (2021).

4. Budd Haeberlein, S., et al. Two Randomized Phase 3 Studies of Aducanumab in Early Alzheimer’s Disease. J Prev Alzheimers Dis 9, 197–210 (2022).

5. Honig, L. S. et al. Trial of Solanezumab for Mild Dementia Due to Alzheimer’s Disease. N Engl J Med 378, 321–330 (2018).

6. Jankowsky, J. L. et al. Mutant presenilins specifically elevate the levels of the 42 residue β-amyloid peptide in vivo: evidence for augmentation of a 42-specific γ secretase. Human Molecular Genetics 13, 159–170 (2004).

7. Jankowsky, J. L. et al. Co-expression of multiple transgenes in mouse CNS: a comparison of strategies. Biomol Eng 17, 157–165 (2001).

8. O’Keefe, J. Place units in the hippocampus of the freely moving rat. Experimental Neurology 51, 78–109 (1976).

9. O’Keefe, J., Burgess, N., Donnett, J. G., Jeffery, K. J. & Maguire, E. A. Place cells, navigational accuracy, and the human hippocampus. Philos Trans R Soc Lond B Biol Sci 353, 1333–1340 (1998).

10. Cacucci, F., Yi, M., Wills, T. J., Chapman, P. & O’Keefe, J. Place cell firing correlates with memory deficits and amyloid plaque burden in Tg2576 Alzheimer mouse model. Proceedings of the National Academy of Sciences 105, 7863–7868 (2008).

11. Mably, A. J., Gereke, B. J., Jones, D. T. & Colgin, L. L. Impairments in spatial representations and rhythmic coordination of place cells in the 3xTg mouse model of Alzheimer’s disease. Hippocampus 27, 378–392 (2017).

12. Belluscio, M. A., Mizuseki, K., Schmidt, R., Kempter, R. & Buzsaki, G. Cross-Frequency Phase-Phase Coupling between Theta and Gamma Oscillations in the Hippocampus. Journal of Neuroscience 32, 423–435 (2012).

13. Colgin, L. L. Rhythms of the hippocampal network. Nat Rev Neurosci 17, 239–249 (2016).

14. Jacobson, T. K. et al. Hippocampal theta, gamma, and theta-gamma coupling: effects of aging, environmental change, and cholinergic activation. Journal of Neurophysiology 109, 1852–1865 (2013).

15. Kendrick, K. M. et al. Learning alters theta amplitude, theta-gamma coupling and neuronal synchronization in inferotemporal cortex. BMC Neuroscience 12, 55 (2011).

16. Lisman, J. E. & Jensen, O. The Theta-Gamma Neural Code. Neuron 77, 1002–1016 (2013).

17. Tamura, M., Spellman, T. J., Rosen, A. M., Gogos, J. A. & Gordon, J. A. Hippocampal-prefrontal theta-gamma coupling during performance of a spatial working memory task. Nat Commun 8, 2182 (2017).

18. Tort, A. B. L., Komorowski, R. W., Manns, J. R., Kopell, N. J. & Eichenbaum, H. Theta– gamma coupling increases during the learning of item–context associations. Proc. Natl. Acad. Sci. U.S.A. 106, 20942–20947 (2009).

19. Vivekananda, U. et al. Theta power and theta-gamma coupling support long-term spatial memory retrieval. Hippocampus 31, 213–220 (2021).

20. Zhou, H. et al. Disruption of hippocampal neuronal circuit function depends upon behavioral state in the APP/PS1 mouse model of Alzheimer’s disease. Sci Rep 12, 21022 (2022).

21. Prince, S. M. et al. Alzheimer’s pathology causes impaired inhibitory connections and reactivation of spatial codes during spatial navigation. Cell Rep 35, 109008 (2021).

22. Zhurakovskaya, E. et al. Impaired hippocampal-cortical coupling but preserved local synchrony during sleep in APP/PS1 mice modeling Alzheimer’s disease. Sci Rep 9, 5380 (2019).

23. Buzsáki, G. Theta Oscillations in the Hippocampus. Neuron 33, 325–340 (2002).

24. Colgin, L. L. Theta–gamma coupling in the entorhinal–hippocampal system. Current Opinion in Neurobiology 31, 45–50 (2015).

25. O’Brien, R. J. & Wong, P. C. Amyloid precursor protein processing and Alzheimer’s disease. Annu Rev Neurosci 34, 185–204 (2011).

26. Arbel-Ornath, M. et al. Soluble oligomeric amyloid-β induces calcium dyshomeostasis that precedes synapse loss in the living mouse brain. Molecular Neurodegeneration 12, 27 (2017).

27. Cleary, J. P. et al. Natural oligomers of the amyloid-β protein specifically disrupt cognitive function. Nat Neurosci 8, 79–84 (2005).

28. Cline, E. N., Bicca, M. A., Viola, K. L. & Klein, W. L. The Amyloid-β Oligomer Hypothesis: Beginning of the Third Decade. JAD 64, S567–S610 (2018).

29. Ding, Y. et al. Amyloid Beta Oligomers Target to Extracellular and Intracellular Neuronal Synaptic Proteins in Alzheimer’s Disease. Frontiers in Neurology 10, (2019).

30. Faucher, P., Mons, N., Micheau, J., Louis, C. & Beracochea, D. J. Hippocampal Injections of Oligomeric Amyloid β-peptide (1–42) Induce Selective Working Memory Deficits and Long-lasting Alterations of ERK Signaling Pathway. Frontiers in Aging Neuroscience 7, (2016).

31. Ferreira, S. T., Lourenco, M. V., Oliveira, M. M. & De Felice, F. G. Soluble amyloid-β oligomers as synaptotoxins leading to cognitive impairment in Alzheimer’s disease. Front Cell Neurosci 9, 191 (2015).

32. Nicole, O. et al. Soluble amyloid beta oligomers block the learning-induced increase in hippocampal sharp wave-ripple rate and impair spatial memory formation. Sci Rep 6, 22728 (2016).

33. Özcan, G. G., Lim, S., Leighton, P. L., Allison, W. T. & Rihel, J. Sleep is bi-directionally modified by amyloid beta oligomers. eLife 9, e53995 (2020).

34. Shankar, G. M. et al. Amyloid-β protein dimers isolated directly from Alzheimer’s brains impair synaptic plasticity and memory. Nat Med 14, 837–842 (2008).

35. Malinow, R. & Malenka, R. C. AMPA receptor trafficking and synaptic plasticity. Annu Rev Neurosci 25, 103–126 (2002).

36. Yang, T., Li, S., Xu, H., Walsh, D. M. & Selkoe, D. J. Large Soluble Oligomers of Amyloid β-Protein from Alzheimer Brain Are Far Less Neuroactive Than the Smaller Oligomers to Which They Dissociate. J Neurosci 37, 152–163 (2017).

37. Busche, M. A. et al. Critical role of soluble amyloid-β for early hippocampal hyperactivity in a mouse model of Alzheimer’s disease. Proc. Natl. Acad. Sci. U.S.A. 109, 8740–8745 (2012).

38. Busche, M. A. & Konnerth, A. Neuronal hyperactivity – A key defect in Alzheimer’s disease? BioEssays 37, 624–632 (2015).

39. Zarhin, D. et al. Disrupted neural correlates of anesthesia and sleep reveal early circuit dysfunctions in Alzheimer models. Cell Reports 38, 110268 (2022).

40. Garcia-Alloza, M. et al. Characterization of amyloid deposition in the APPswe/PS1dE9 mouse model of Alzheimer disease. Neurobiol Dis 24, 516–524 (2006).

41. Takeda, S. et al. Brain interstitial oligomeric amyloid β increases with age and is resistant to clearance from brain in a mouse model of Alzheimer’s disease. FASEB J 27, 3239–3248 (2013).

42. Zhang, L. et al. Dynamic Changes in the Levels of Amyloid-β42 Species in the Brain and Periphery of APP/PS1 Mice and Their Significance for Alzheimer’s Disease. Front Mol Neurosci 14, 723317 (2021).

43. Webster, S. J., Bachstetter, A. D. & Van Eldik, L. J. Comprehensive behavioral characterization of an APP/PS-1 double knock-in mouse model of Alzheimer’s disease. Alzheimers Res Ther 5, 28 (2013).

44. Knierim, J. J., Kudrimoti, H. S. & McNaughton, B. L. Place cells, head direction cells, and the learning of landmark stability. J Neurosci 15, 1648–1659 (1995).

45. Jun, H. et al. Disrupted Place Cell Remapping and Impaired Grid Cells in a Knockin Model of Alzheimer’s Disease. Neuron 107, 1095–1112.e6 (2020).

46. Ego-Stengel, V. & Wilson, M. A. Disruption of ripple-associated hippocampal activity during rest impairs spatial learning in the rat. Hippocampus 20, 1–10 (2010).

47. Ólafsdóttir, H. F., Bush, D. & Barry, C. The Role of Hippocampal Replay in Memory and Planning. Curr Biol 28, R37–R50 (2018).

48. Wilson, M. A. & McNaughton, B. L. Reactivation of Hippocampal Ensemble Memories During Sleep. Science 265, 676–679 (1994).

49. Siapas, A. G. & Wilson, M. A. Coordinated Interactions between Hippocampal Ripples and Cortical Spindles during Slow-Wave Sleep. Neuron 21, 1123–1128 (1998).

50. Born, J., Rasch, B. & Gais, S. Sleep to Remember. Neuroscientist 12, 410–424 (2006).

51. Bazzigaluppi, P. et al. Early-stage attenuation of phase-amplitude coupling in the hippocampus and medial prefrontal cortex in a transgenic rat model of Alzheimer’s disease. Journal of Neurochemistry 144, 669–679 (2018).

52. Etter, G. et al. Optogenetic gamma stimulation rescues memory impairments in an Alzheimer’s disease mouse model. Nat Commun 10, 5322 (2019).

53. Gurevicius, K., Lipponen, A. & Tanila, H. Increased Cortical and Thalamic Excitability in Freely Moving APPswe/PS1dE9 Mice Modeling Epileptic Activity Associated with Alzheimer’s Disease. Cerebral Cortex 23, 1148–1158 (2013).

54. Stoiljkovic, M., Kelley, C., Stutz, B., Horvath, T. L. & Hajós, M. Altered Cortical and Hippocampal Excitability in TgF344-AD Rats Modeling Alzheimer’s Disease Pathology. Cerebral Cortex 29, 2716–2727 (2019).

55. Rodriguez-Larios, J. & Alaerts, K. Tracking Transient Changes in the Neural Frequency Architecture: Harmonic Relationships between Theta and Alpha Peaks Facilitate Cognitive Performance. J Neurosci 39, 6291–6298 (2019).

56. Bandarabadi, M. et al. Dynamic modulation of theta-gamma coupling during rapid eye movement sleep. Sleep 42, zsz182 (2019).

57. Montgomery, S. M., Sirota, A. & Buzsáki, G. Theta and Gamma Coordination of Hippocampal Networks during Waking and Rapid Eye Movement Sleep. J. Neurosci. 28, 6731–6741 (2008).

58. Joo, H. R. & Frank, L. M. The hippocampal sharp wave-ripple in memory retrieval for immediate use and consolidation. Nat Rev Neurosci 19, 744–757 (2018).

59. Girardeau, G., Cei, A. & Zugaro, M. Learning-Induced Plasticity Regulates Hippocampal Sharp Wave-Ripple Drive. J. Neurosci. 34, 5176–5183 (2014).

60. Atherton, L. A., Dupret, D. & Mellor, J. R. Memory trace replay: the shaping of memory consolidation by neuromodulation. Trends Neurosci 38, 560–570 (2015).

61. Louie, K. & Wilson, M. A. Temporally Structured Replay of Awake Hippocampal Ensemble Activity during Rapid Eye Movement Sleep. Neuron 29, 145–156 (2001).

62. Kurucu, H. et al. Inhibitory synapse loss and accumulation of amyloid beta in inhibitory presynaptic terminals in Alzheimer’s disease. European Journal of Neurology 29, 1311–1323 (2022).

63. Tang, W. & Jadhav, S. P. Multiple-Timescale Representations of Space: Linking Memory to Navigation. Annu Rev Neurosci 45, 1–21 (2022).

64. Lisman, J. The theta/gamma discrete phase code occuring during the hippocampal phase precession may be a more general brain coding scheme. Hippocampus 15, 913–922 (2005).

65. Skaggs, W. E., McNaughton, B. L., Wilson, M. A. & Barnes, C. A. Theta phase precession in hippocampal neuronal populations and the compression of temporal sequences. Hippocampus 6, 149–172 (1996).

66. Lonnemann, N., Korte, M. & Hosseini, S. Repeated performance of spatial memory tasks ameliorates cognitive decline in APP/PS1 mice. Behavioural Brain Research 438, 114218 (2023).

67. Kilgore, M. et al. Inhibitors of Class 1 Histone Deacetylases Reverse Contextual Memory Deficits in a Mouse Model of Alzheimer’s Disease. Neuropsychopharmacology 35, 870–880 (2010).

68. Medeiros, R., Baglietto-Vargas, D. & LaFerla, F. M. The role of tau in Alzheimer’s disease and related disorders. CNS Neurosci Ther 17, 514–524 (2011).

69. Niikura, T., Tajima, H. & Kita, Y. Neuronal Cell Death in Alzheimer’s Disease and a Neuroprotective Factor, Humanin. Curr Neuropharmacol 4, 139–147 (2006).

70. Lambert, M. P. et al. Diffusible, nonfibrillar ligands derived from Abeta1-42 are potent central nervous system neurotoxins. Proc Natl Acad Sci U S A 95, 6448–6453 (1998).

71. Hampel, H. et al. The Amyloid-β Pathway in Alzheimer’s Disease. Mol Psychiatry 26, 5481– 5503 (2021).

72. Chow, V. W., Mattson, M. P., Wong, P. C. & Gleichmann, M. An Overview of APP Processing Enzymes and Products. Neuromolecular Med 12, 1–12 (2010).

73. Bukhari, H. et al. Small things matter: Implications of APP intracellular domain AICD nuclear signaling in the progression and pathogenesis of Alzheimer’s disease. Prog Neurobiol 156, 189–213 (2017).

74. Kreis, A. et al. Overexpression of wild-type human amyloid precursor protein alters GABAergic transmission. Sci Rep 11, 17600 (2021).

75. Do, H. N., Devkota, S., Bhattarai, A., Wolfe, M. S. & Miao, Y. Effects of presenilin-1 familial Alzheimer’s disease mutations on γ-secretase activation for cleavage of amyloid precursor protein. Commun Biol 6, 1–14 (2023).

76. Soto-Faguás, C. M., Sanchez-Molina, P. & Saura, C. A. Loss of presenilin function enhances tau phosphorylation and aggregation in mice. Acta Neuropathol Commun 9, 162 (2021).

77. Kelleher, R. J. & Shen, J. Presenilin-1 mutations and Alzheimer’s disease. Proc Natl Acad Sci U S A 114, 629–631 (2017).

78. Woodruff, G. et al. The Presenilin-1 ΔE9 mutation results in reduced γ-secretase activity, but not total loss of PS1 function, in isogenic human stem cells. Cell Rep 5, 10.1016/j.celrep.2013.10.018 (2013).

79. Bagaria, J., Bagyinszky, E. & An, S. S. A. Genetics, Functions, and Clinical Impact of Presenilin-1 (PSEN1) Gene. Int J Mol Sci 23, 10970 (2022).

80. Zhou, H. et al. Cholinergic modulation of hippocampal calcium activity across the sleep-wake cycle. eLife 8, e39777 (2019).

81. Li, X. et al. Sex differences between APPswePS1dE9 mice in A-beta accumulation and pancreatic islet function during the development of Alzheimer’s disease. Lab Anim 50, 275– 285 (2016).

82. Zhou, P. et al. Efficient and accurate extraction of in vivo calcium signals from microendoscopic video data. eLife 7, e28728 (2018).

83. Lerdkrai, C. et al. Intracellular Ca ^2+^ stores control in vivo neuronal hyperactivity in a mouse model of Alzheimer’s disease. Proc. Natl. Acad. Sci. U.S.A. 115, (2018).

84. Markus, E. J., Barnes, C. A., McNaughton, B. L., Gladden, V. L. & Skaggs, W. E. Spatial information content and reliability of hippocampal CA1 neurons: Effects of visual input. Hippocampus 4, 410–421 (1994).

85. Go, M. A. et al. Place Cells in Head-Fixed Mice Navigating a Floating Real-World Environment. Frontiers in Cellular Neuroscience 15, (2021).

86. Bokil, H., Andrews, P., Kulkarni, J. E., Mehta, S. & Mitra, P. P. Chronux: A platform for analyzing neural signals. Journal of Neuroscience Methods 192, 146–151 (2010).

87. Onslow, A. C. E., Bogacz, R. & Jones, M. W. Quantifying phase–amplitude coupling in neuronal network oscillations. Progress in Biophysics and Molecular Biology 105, 49–57 (2011).

88. Osipova, D., Hermes, D. & Jensen, O. Gamma Power Is Phase-Locked to Posterior Alpha Activity. PLoS ONE 3, e3990 (2008).

