## Supplementary figures and tables for "Impaired hippocampal circuit function underlying memory encoding and consolidation precede robust Aβ deposition in a mouse model of Alzheimer’s disease"

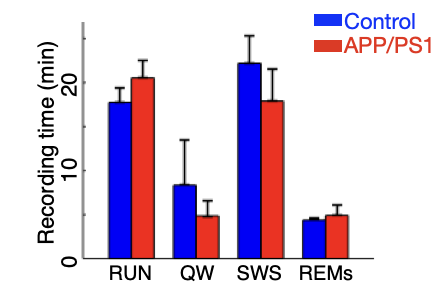

***Supplementary Figure S1. Recording length of each state is comparable between genotypes. p>0.05, 2-sided t-tests.***

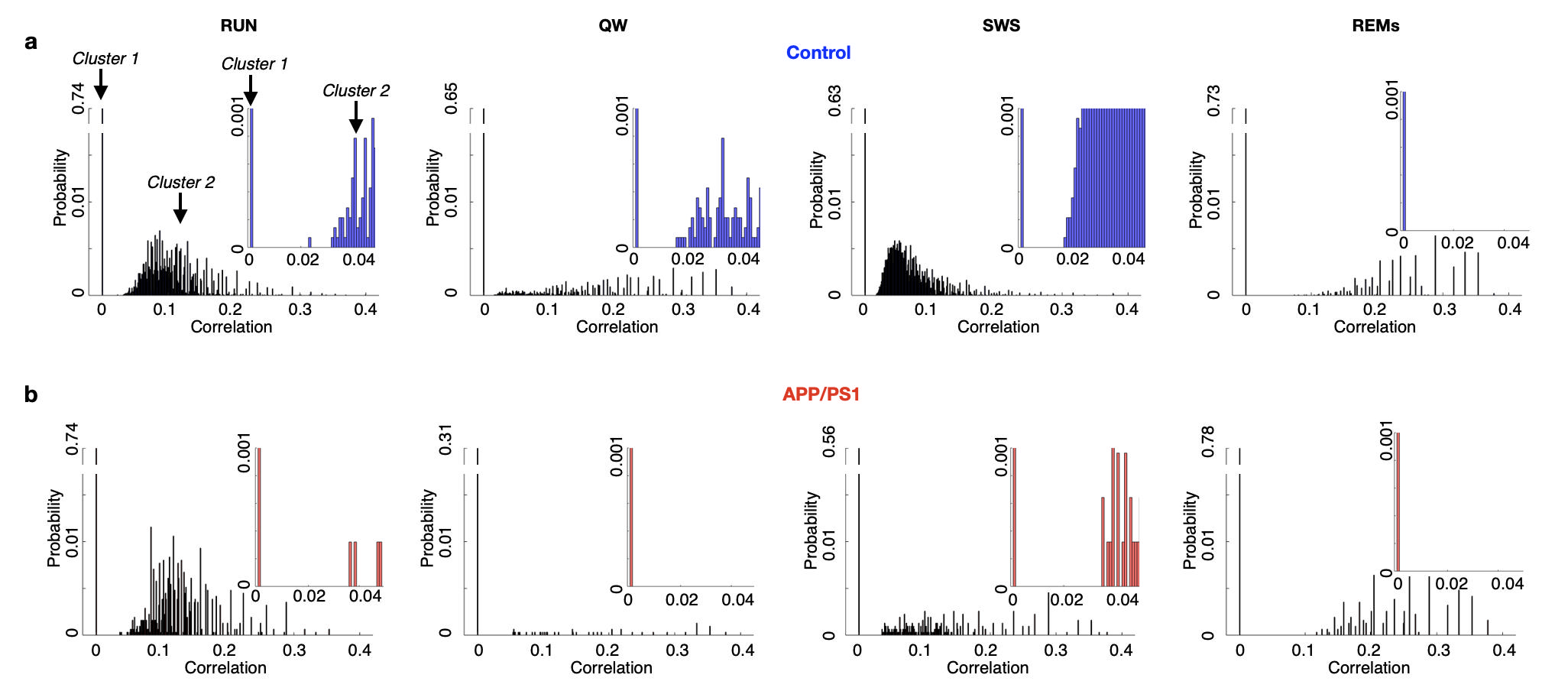
***Supplementary Figure S2. Distribution of place cell pairwise cross correlations at zero lag.*** *In each behavioral state of* ***a)*** *control and* ***b)*** *APP/PS1 mice, two clusters were observed: the 1^st^ bin (bin size = 0.001, cluster 1) and the rest of the distribution (cluster 2). We define cluster 2 as coactive. A threshold of coactivity of 0.002 adequately distinguished these clusters. Insert: zoom in 0 to 0.05 to visualize the gap between the clusters. Control n=5 mice, 395 place cells; APP/PS1 n=6 mice, 237 place cells.*

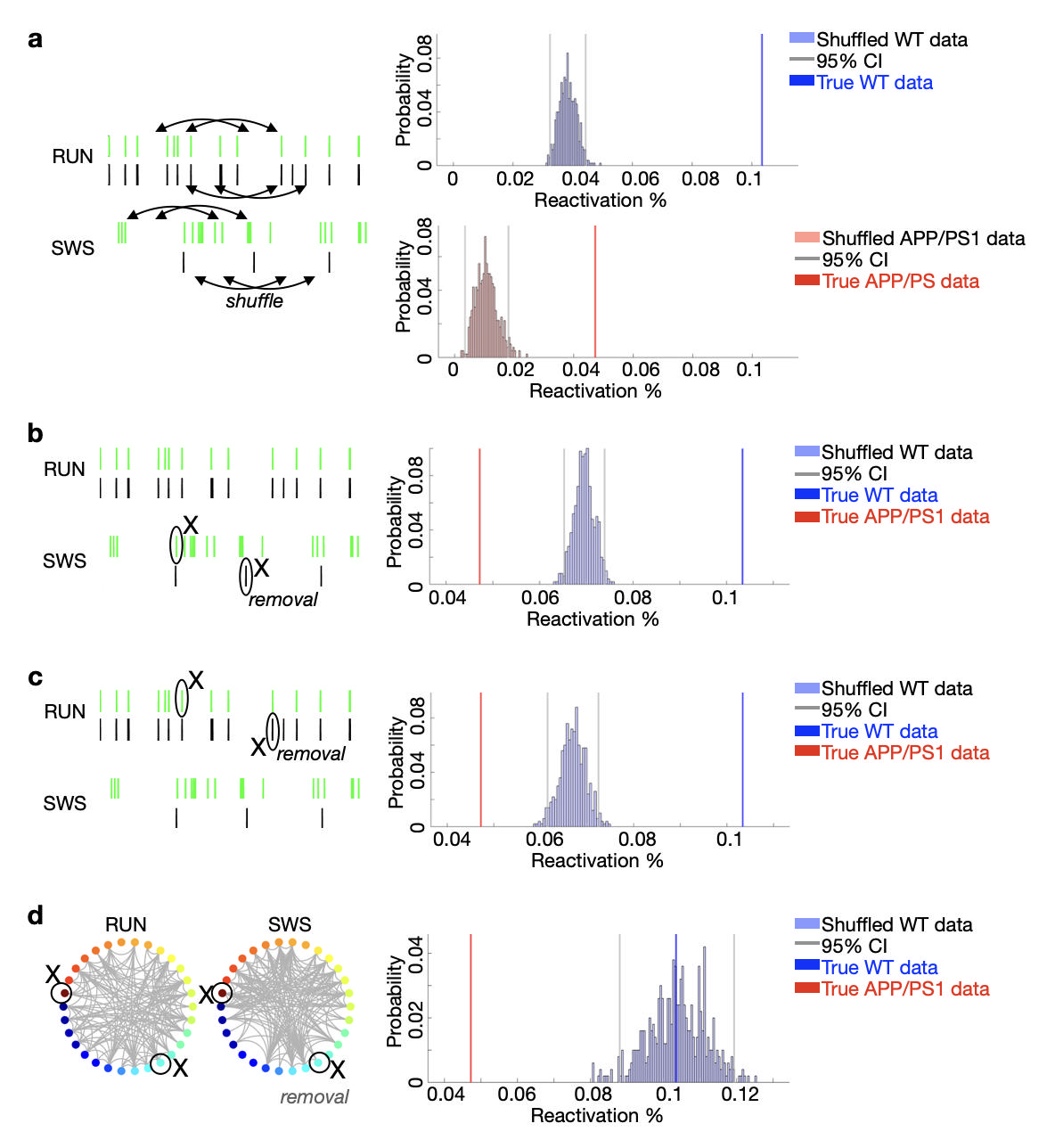

***Supplementary Figure S3. Reduced APP/PS1 place cell pairwise reactivation in SWS persists after shuffling. a)*** *In both control (WT) and APP/PS1 mice, the proportion of place cell pairwise reactivation was significantly greater than the proportion predicted by the distribution of cross-correlations computed using shuffled RUN and SWS spike times (N=500 shuffles).* ***b)*** *After randomly removing spikes from SWS in control animals to match the SWS mean firing rate in APP/PS1, the distribution of cross-correlations of shuffled downsampled SWS and RUN spike times of control mice (N=500) remained higher than place cell pairwise reactivation in APP/PS1 mice.* ***c)*** *After randomly removing spikes from RUN in control animals to match the RUN firing rate in APP/PS1, the distribution of cross-correlations of shuffled SWS and downsampled RUN spike times of control mice (N=500) remained higher than place cell pairwise reactivation in APP/PS1 mice**.* ***d)*** *After randomly removing place cells from each control animal to match the average place cell number in APP/PS1, the distribution of cross-correlations of SWS and RUN spike times of control mice (N=500) remained higher than place cell pairwise reactivation in APP/PS1 mice.*

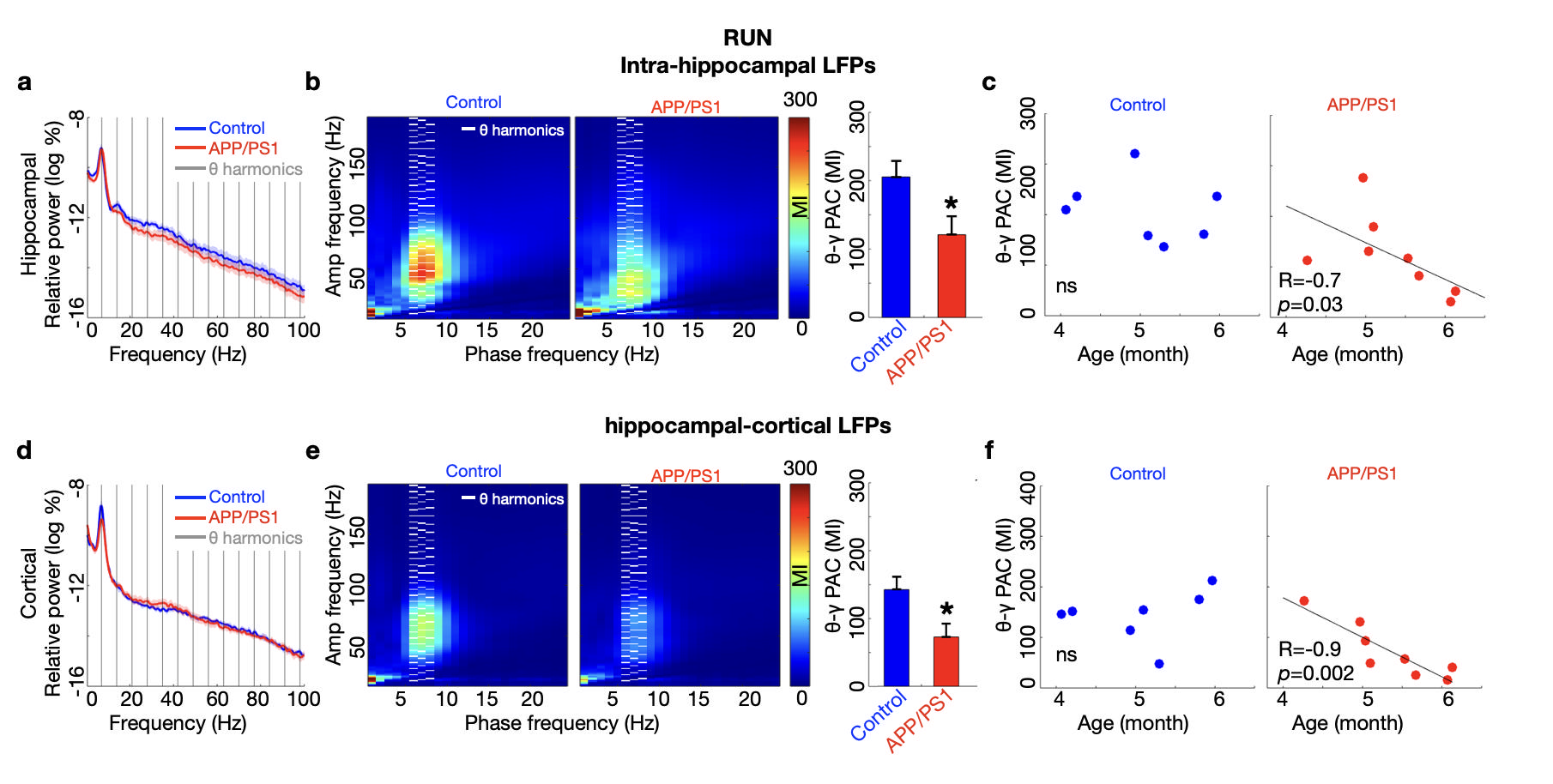
***Supplementary Figure S4. Impairment of online theta-gamma phase amplitude coupling (PAC) and its decrease with age in young APP/PS1 mice persists after theta harmonics are excluded from the PAC measurement. a)*** *In RUN, hippocampal theta and gamma power in APP/PS1 and control mice were comparable. Grey lines: theta harmonics. After subtracting the PAC at theta harmonic frequencies (multiples of theta peak frequency 7Hz):* ***b)*** *Intra-hippocampal theta (6-8Hz) - gamma (40-80Hz) PAC was reduced in APP/PS1 compared to control mice. White lines: theta harmonics.* ***c)*** *Intrahippocampal theta-gamma PAC was negatively correlated with the age of APP/PS1 mice but not control mice.* ***d)*** *Cortical gamma power was comparable between genotypes. After subtracting the PAC at theta harmonic frequencies:* ***e)*** *PAC of hippocampal theta (6-8Hz) - cortical gamma (40-80Hz) was reduced in APP/PS1 compared to control mice.* ***f)*** *The reduction of hippocampal theta- cortical gamma PAC was negatively correlated with age in APP/PS1 but not control mice. *p < 0.05; 2-sided t-tests, Pearson correlation. Control n=7 mice; APP/PS1 n=8 mice. Data are represented as mean ± S.E.M.*

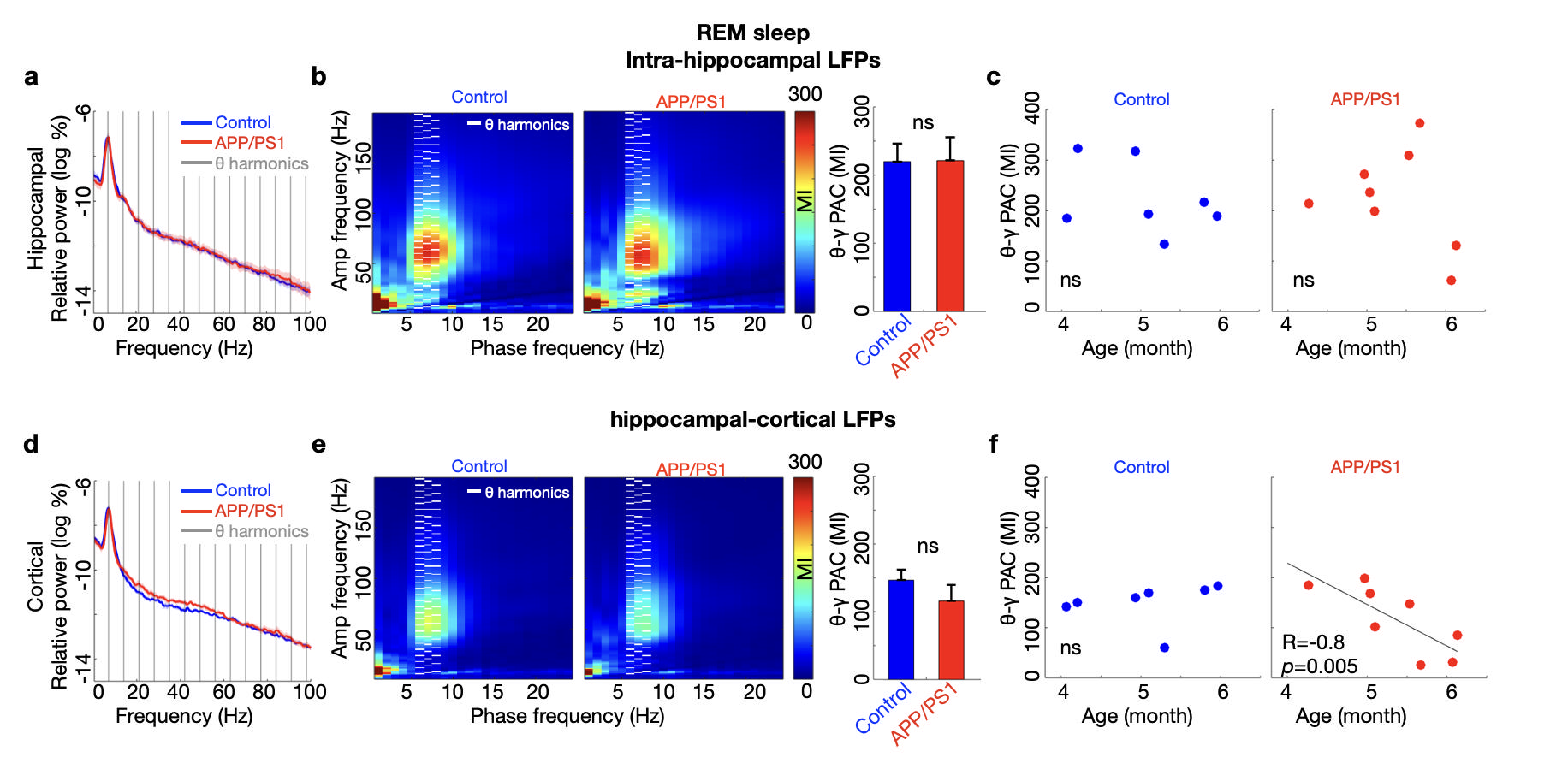
***Supplementary Figure S5. Preservation of REM sleep theta-gamma phase amplitude coupling (PAC) and its decrease with age in young APP/PS1 mice after theta harmonics are excluded from the PAC measurement. a)*** *In REM sleep, hippocampal theta and gamma power in APP/PS1 and control mice were comparable. Grey lines: theta harmonics. After subtracting the PAC at theta harmonic frequencies (multiples of theta peak frequency 7Hz:* ***b)*** *Intra-hippocampal theta (6-8Hz) - gamma (40-80Hz) PAC was intact in APP/PS1 mice. White lines: theta harmonics.* ***c)*** *Intrahippocampal theta-gamma PAC was not correlated with age in either genotype.* ***d)*** *Cortical gamma power was comparable between genotypes. After subtracting the PAC at theta harmonic frequencies (multiples of theta peak frequency 7Hz):* ***e)*** *PAC of hippocampal theta (6-8Hz) - cortical gamma (40-80Hz) was intact in APP/PS1 compared to control mice.* ***f)*** *The reduction of hippocampal theta- cortical gamma PAC was negatively correlated with age in APP/PS1 but not control mice. 2-sided t-tests, Pearson correlation. Control n=7 mice; APP/PS1 n=8 mice. Data are represented as mean ± S.E.M.*

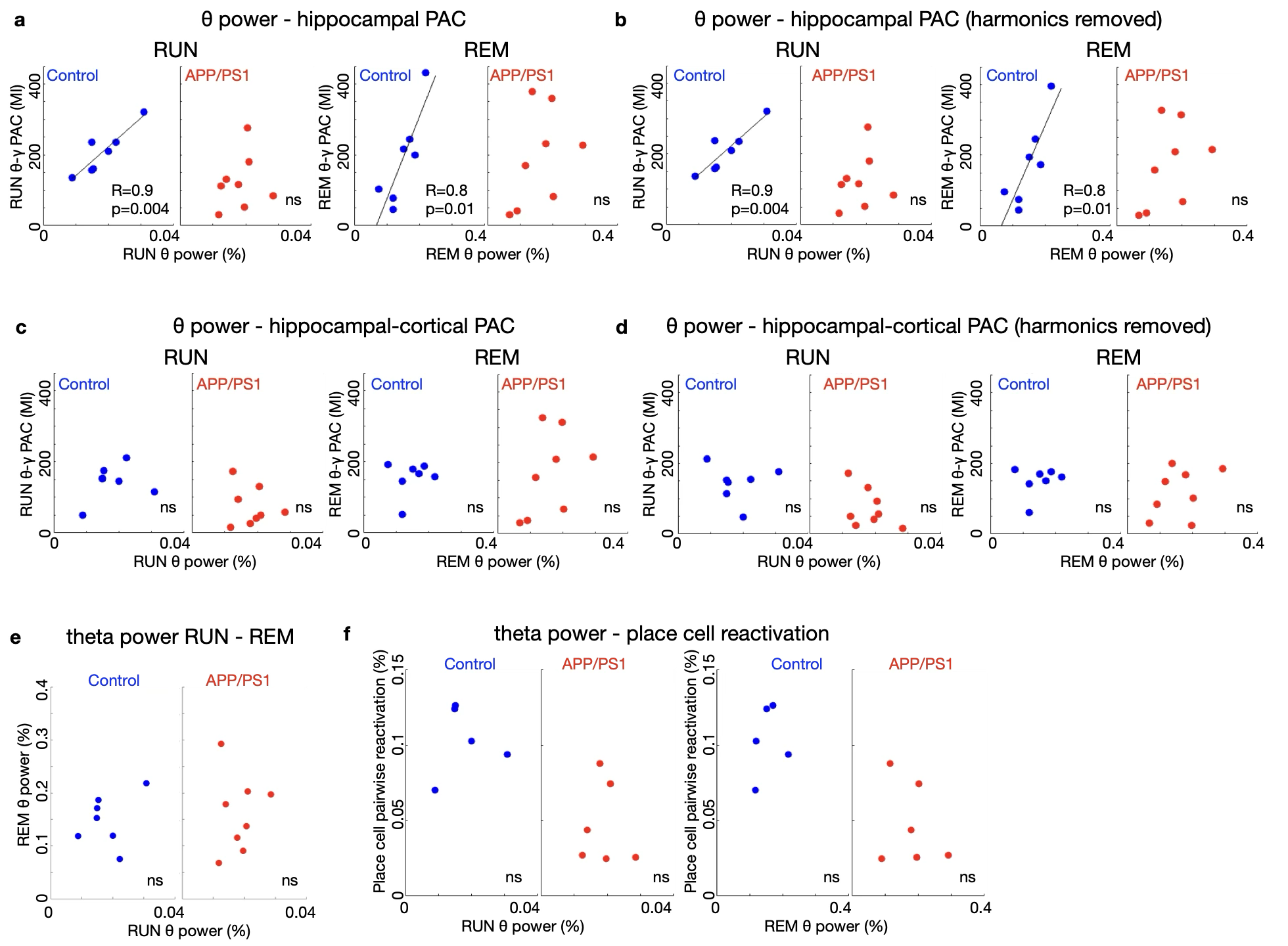

***Supplementary Figure S6. Theta power is correlated with theta-gamma phase amplitude coupling in Control mice but not in APP/PS1 mice.*** *a) Hippocampal theta power were significantly correlated with hippocampal theta-gamma phase-amplitude coupling (PAC) in RUN and REM sleep in control mice, but not in APP/PS1 mice. b) This result persisted when theta harmonics were removed from the PAC analysis. c) Hippocampal theta power was not significantly correlated with hippocampal theta- cortical gamma in RUN or REM sleep in control or APP/PS1 mice. d) This result persisted when theta harmonics were removed from the PAC analysis. e) Theta power in RUN was not significantly correlated with theta power in REM sleep in either genotype. f) Theta power in RUN was not significantly correlated with place cell pairwise reactivation in slow wave sleep in either genotype. Pearson correlation. Control n=7 mice; APP/PS1 n=8 mice.*

Supplementary Table 1. Place cell calcium rates ANOVA and post-hoc results

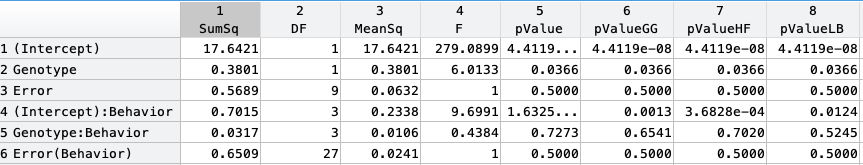

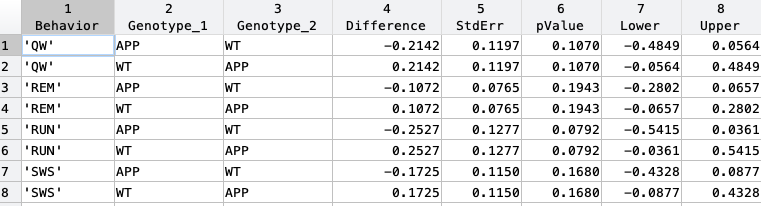

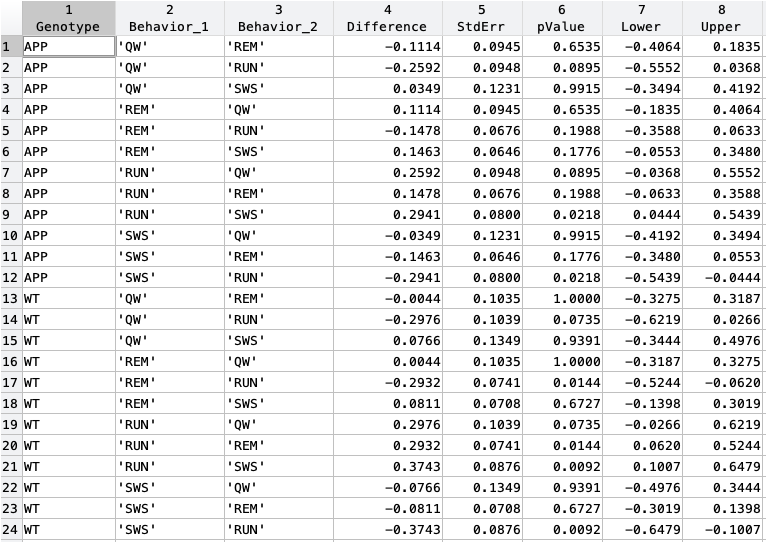

Supplementary Table 2. Non-place cell calcium rates ANOVA and post-hoc results

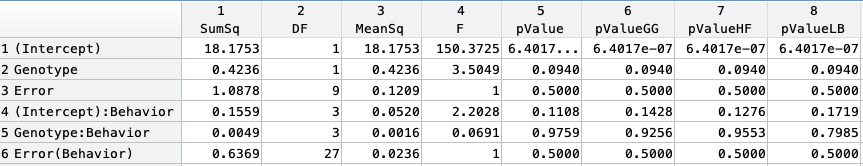

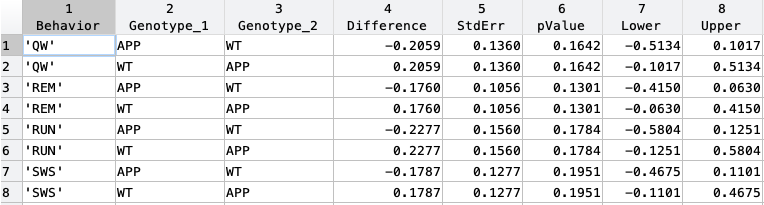

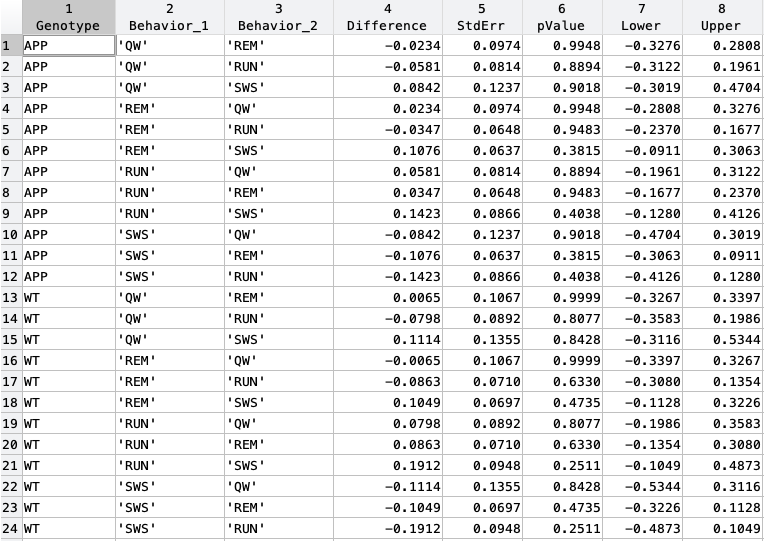

Supplementary Table 3. Place cell reactivation ANOVA and post-hoc results

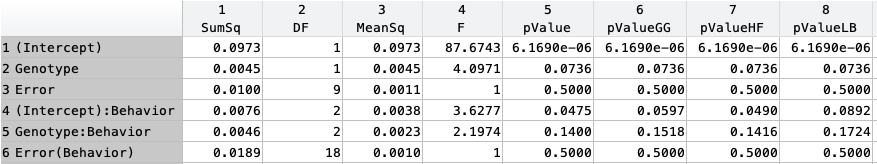

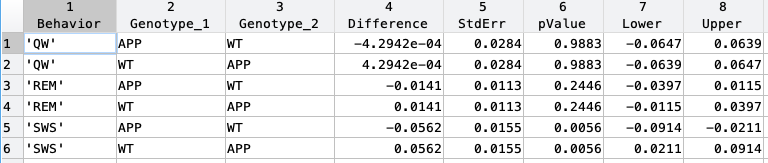

Supplementary Table 4. RUN cell-LFP synchronization ANOVA and post-hoc results

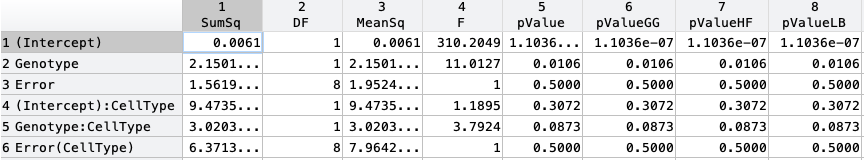

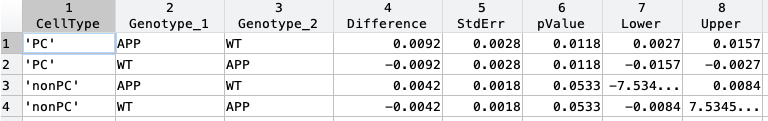

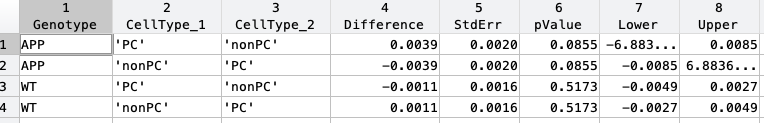

Supplementary Table 5. REM cell-LFP synchronization ANOVA and post-hoc results

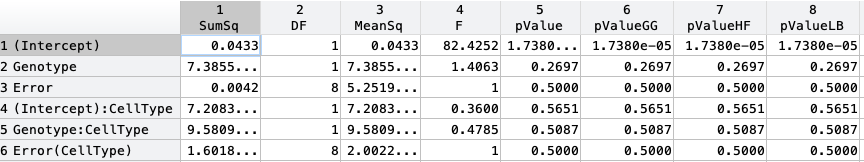

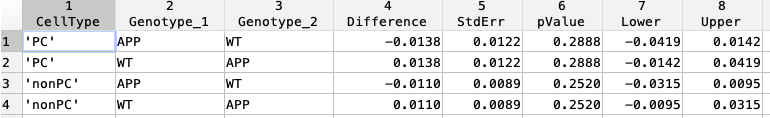

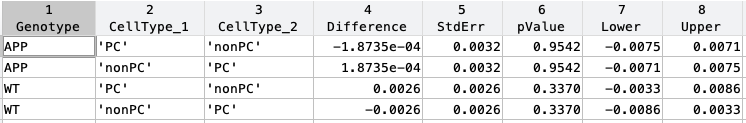
